# Understanding the kinetics of macrophage uptake and the metabolic fate of iron-carbohydrate complexes used for iron deficiency anemia treatment

**DOI:** 10.1101/2025.07.01.662524

**Authors:** Stephanie Eitner, Vera M. Kissling, Alexandra Rippl, Leonard Krupnik, Justus J. Bürgi, Stefanie Gächter, Katharina Hast-Sribike, Alexander Gogos, Davide Bottone, Jonas Bossart, Rafael Riudavets Puig, Reinaldo Digigow, Amy E. Alston, Beat Flühmann, Peter Wick, Vanesa Ayala-Nunez

## Abstract

Iron-carbohydrate complexes (ICCs) are widely used nanomedicines to treat iron deficiency anemia, yet their intracellular fate and the mechanisms of action underlying their differences in treatment outcomes remain poorly understood. Here, we thus performed a comprehensive dynamic characterization of two structurally distinct ICCs ‒ iron sucrose (IS) and ferric carboxymaltose (FCM) ‒ in primary human macrophages, key cells to the iron metabolism. By employing innovative correlative microscopy techniques, elemental analysis, and *in vitro* pharmacokinetic profiling, we demonstrate that the uptake, intracellular trafficking, and biodegradation of ICCs depend on their physicochemical properties. Specifically, IS is rapidly internalized and processed within endolysosomes, resulting in fast iron release and transient cytotoxicity. Conversely, FCM is sequestered in enlarged endosomes for an extended time before its biodegradation, a phenomenon we term the *Hamster Effect*, which leads to slower, more sustainable iron release. These results provide unprecedented insights into the metabolic fate of ICCs, enhancing our understanding of their different pharmacokinetic and pharmacodynamic profiles *in vivo*.

---

Iron deficiency and iron deficiency anemia are global public health problems that affect over 1 billion people worldwide.^1^ Intravenous iron-carbohydrate complexes (ICCs) are nanomedicines that are widely used to treat iron deficiency and iron deficiency anemia when oral iron supplementation is not feasible or intolerable to the patient.^2,3^ The approved ICCs differ with regard to their polynuclear iron core crystallinity, size, surface chemistry and other physicochemical characteristics (PCCs).^4^ In patients, each ICC exhibits different pharmacokinetic (PK) and pharmacodynamic (PD) profiles that are defined by their diverse PCCs.^5^ To understand how the PCCs of these nanomedicines define their PK/PD in patients, we need to unravel the mechanisms at the nano-bio interface, i.e. their dynamic interaction profiles at the cellular level. The prevailing hypothesis for ICC handling after injection has been that the ICCs are initially internalized by macrophages from the reticuloendothelial system, predominantly in the liver and spleen.^6^ Yet, despite more than seven decades of successful clinical use with several ICCs available on the market, a complete understanding of the nano-bio interface in macrophages ‒ key players in the body’s iron metabolism ‒ is still lacking. This is likely in part due to the lack of advanced characterization and imaging methodologies available at the time of the ICCs’ development.

In this study, we thus investigated the dynamics of two structurally distinct ICCs ‒ iron sucrose (IS) and ferric carboxymaltose (FCM) ‒ within primary human macrophages, utilizing an innovative and correlative combination of advanced iron staining techniques, fluorescence and electron microscopy, small-angle X-ray scattering (SAXS), elemental iron analysis, and several cell-based assays. Our main aim was to unravel how different ICCs are taken up by macrophages, converted from their nanoparticle form into bioavailable iron (i.e. intracellular biodegradation, storage and mobilization), and finally integrated into the iron metabolism of the cell. To this end, we employed an experimental approach based on the recent understanding of the PCCs of ICCs by our group^7^ and existing clinical knowledge.^5^ The findings of this multi-modal, systematic investigation may be further applied to other metal oxide nanoparticles to enhance understanding of their *in vivo* fate using *in vitro* techniques.

## Results

### What are the cells seeing? Characterization of ICCs in cell culture medium

The limited mechanistic understanding of ICCs stems primarily from their highly complex physicochemical properties,^8^ which render their investigation in biological systems particularly challenging. Our recent work unveiled the intricate structure of ICCs in water and saline solution, showing that they are highly polydisperse and composed of a mixture of single nanoparticles (NPs), clusters and agglomerates of varied sizes.^7,9^ Moreover, it has been reported that ICCs can contain a fraction of labile and free iron (IS: 0.69-4.5%, FCM: <0.5%).^10^ Additional complexity is introduced after intravenous administration, when labile iron (LI) can bind to transferrin to form transferrin-bound iron (TBI), or to other serum proteins and molecules (e.g. albumin, citrate) to form non-transferrin bound iron (NTBI). Notably, both NP-complexed iron and non-complexed iron can have a biological relevance in patients.^11^

Considering this complex scenario where different iron species co-exist, we first characterized the ICCs in cell culture medium using standard cell culture conditions for NP-exposure of primary human macrophages *in vitro* (RPMI, 10% FCS, 1% Pen-Strep). In bright-field transmission electron microscopy (BF-TEM), IS appeared mainly as separate small, spherical nanoparticles (NPs, outlined arrow) with occasional clusters of these NPs and agglomerates (black arrows) in water, while FCM predominantly occurred as ellipsoidal clusters of nanoparticles (black arrows) (Fig.1a, Table 1). In cell culture medium (Fig.1b), the separate IS NPs interacted with the medium components, leading to a higher proportion of strongly agglomerated NPs (black arrows, Table 1) with co-localizing "fuzzy" agglomerations of lower electron contrast (white arrows) that are likely medium proteins. Such particle agglomeration in complete cell culture media has been reported also for other NPs.^12^ Conversely, FCM did not change in cluster or agglomerate size, but showed slightly smaller inter-cluster distances (Fig.1b, black arrows). Unbound medium proteins were visible in the background (white arrows), consistent with previous reports of undetectable interactions between FCM and medium proteins in SAXS.^7^ Iron citrate (IC), used as a non-nanoparticle Fe^3+^ iron control, formed non-NP precipitates in water (Fig.1a-b, black arrows) and large agglomerates with proteins in medium (Fig.1b, white arrows). Agglomeration of IS but not FCM in cell culture medium was confirmed by SAXS (Fig.1c), where FCM showed no significant changes in colloidal stability in RPMI compared to ultra-pure water over time (1 h, 6 h and 24 h), while IS indicated agglomerate formation (black arrows). Our UV-VIS observations (Fig.1d, Suppl. Fig.1) suggest that transient low-affinity interactions between the FCM carbohydrate ligand surface and medium components occur. Unlike IS, these interactions likely would not lead to the formation of FCM-protein agglomerates, which would be detectable by BF-TEM or SAXS, but may rather influence the iron core’s binding strength to the carbohydrate ligand in FCM. We also confirmed the higher content of LI of IS compared to FCM when diluted in cell culture medium with inductively coupled plasma optical emission spectroscopy (ICP-OES) (Fig.1e-f). Notably, the LI fraction did not significantly increase over time, indicating no LI release by the IS NPs after 24 h of incubation (Fig.1e). For FCM, the LI fraction was undetectable, while for the Fe^3+^ iron control IC the LI was expectedly high (Fig.1f). A cell-free Prussian blue assay also showed that IS has a higher fraction of non-NP complexed iron compared to FCM (Fig.1g).

**Figure 1:**
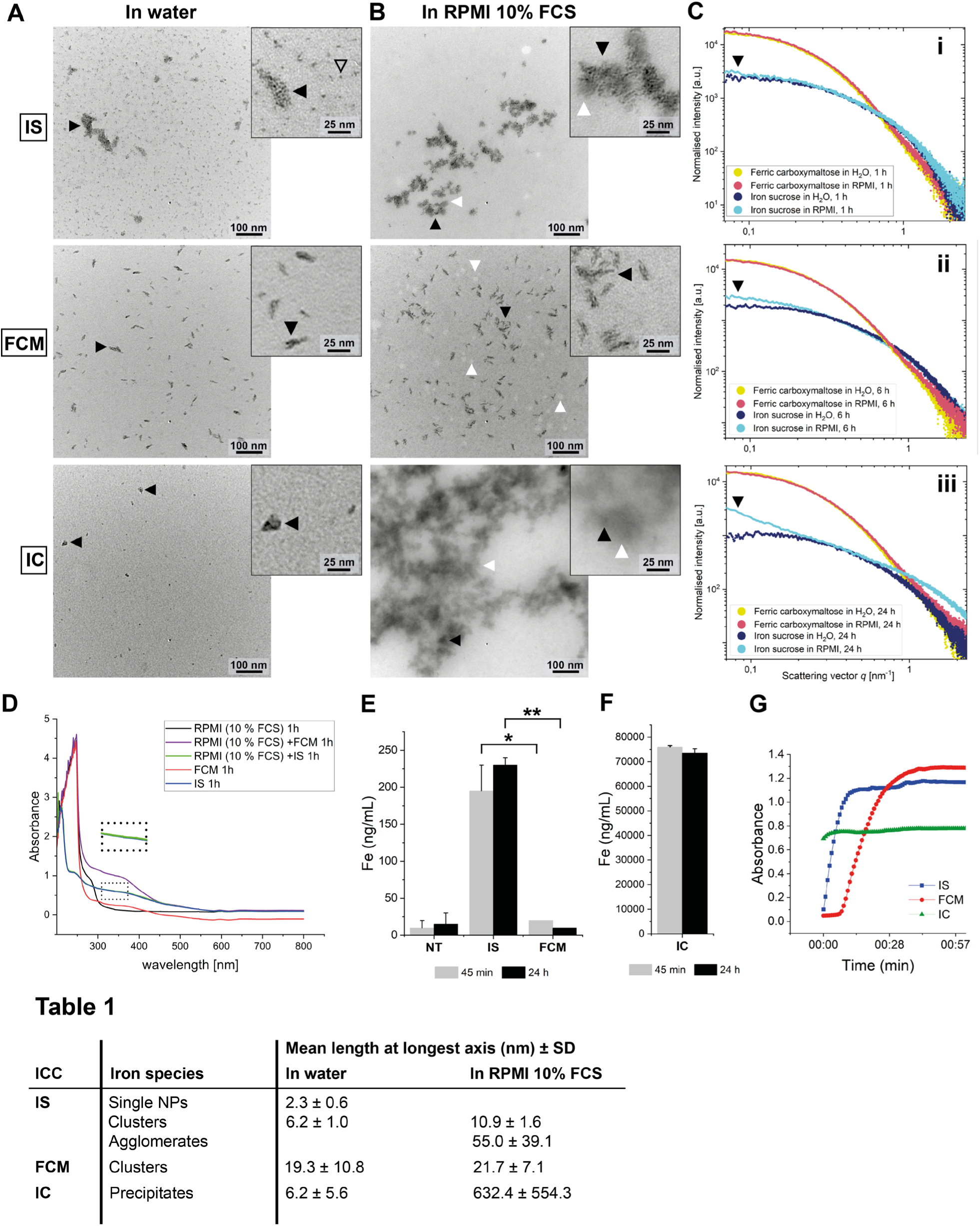
Characterization of iron carbohydrate complexes (ICCs) in cell culture medium. A-B) Representative bright field-TEM images of iron sucrose (IS) and ferric carboxymaltose (FCM) nanoparticles (NPs) as well as the labile Fe^3+^ iron control iron citrate (IC) in water **(A)** and in full cell culture medium (RPMI 10% FCS, 1% Pen-Strep) **(B)** after 24 h incubation at 37 °C, which is a commonly used exposure setting for cell experiments shown later. **A)** Zoom-insets highlight the single spherical NPs visible for IS (black outlined arrow) as well as clusters and agglomerates of IS NPs, the ellipsoidal clusters of FCM NPs and precipitates of IC (black arrows). **B)** Black arrows indicate strong agglomeration of IS NPs and IC precipitates as well as their binding to medium components like proteins (white arrows), which is absent in FCM that showed unbound medium components in the image background (white arrows) and smaller inter-cluster distances without significant size changes of the individual FCM clusters (black arrows, see **Table 1** with mean lengths of iron species at longest axis ± S.D.). Scale bars: 100 nm, 25 nm. **C)** Small-angle X-ray scattering (SAXS) of IS and FCM in water (H_2_O) or RPMI complete medium measured at 1 h **(i)**, 6 h **(ii)** and 24 h **(iii)** of incubation. The curve up-turn (black arrows) of IS in medium compared to water at low *q* indicates clustering of IS NPs, likely with medium components, which is absent in FCM. a.u.: arbitrary units. **D)** UV-VIS measurements of IS and FCM incubated for 1 h with water or complete cell culture medium. The curves of IS 1 h (blue) and IS 1 h in RPMI (10% FCS, green) are overlapping (see dotted square zoom-in). Absorbance: arbitrary units. **E-F)** Quantification of labile iron content in the dilutions of ICCs in cell culture media. A dilution of 1800 µM of iron of IS and FCM was incubated for 45 min and 24 h. Afterwards, they were filtered to separate the labile iron from the complexed iron. Total iron in the flowthrough was quantified with inductively coupled plasma optical emission spectroscopy (ICP-OES). The baseline iron content in the non-treated (NT) control cells stems from iron already present in the medium. The bar graphs represent the mean + S.E.M. of a duplicate. *(0.05 > *P* > 0.01), **(0.01 > *P* > 0.001). **F)** IC was used as a control. **G)** A dynamic Prussian blue assay was performed with IS, FCM and IC diluted in complete cell culture medium. Absorbance (arbitrary units) was measured every 53 s for 1 h.

Taken together, we can conclude that in our experimental conditions, the cells are exposed to a complex mixture of iron species, which is ICC-dependent. In complete cell culture medium, IS likely forms NP-protein agglomerates of different sizes, as well as TBI, NTBI and labile iron. In contrast, the cells treated with FCM probably encounter a mixture of iron species that is stable over time: FCM clusters, less FCM-protein agglomerates, and a very low proportion of non-complexed iron. This difference between IS and FCM can be explained by their carbohydrate ligands: FCM possesses a strongly bound carboxymaltose ligand, contrary to IS with a loosely bound sucrose layer.^7^

### Cell uptake dynamics: complete NPs are internalized *via* endocytosis, a process influenced by the carbohydrate ligand

The complex cell uptake of ICCs is likely driven by the multiple iron species (NP-bound iron, Fe^3+^, Fe^2+^, TBI, NTBI) that are derived from the individual ICC, and their relative concentrations directly related to their physicochemical characteristics.^5,13,14^ A recent pharmacokinetic study demonstrated that all types of iron species are likely influential for the *in vivo* performance and the measured PD/PK profiles of the individual products.^5^

To study the cell uptake dynamics, primary human M2 macrophages were exposed to IS, FCM and IC over the course of 45 min to 48 h, a time frame based on clinical trials conducted with ICCs.^5^ As visible in Fig.2a, the dynamics of uptake measured by ICP-OES were product-dependent: while IS was internalized by the macrophages already after 45 min of exposure and with continuously increased uptake, the internalization of FCM was observed to just begin to increase at 24 h. A faster cell uptake of IS is in accordance with clinical data, where a shorter t_1/2_ was reported.^6^ Conversely, IC’s uptake did not considerably increase over time. A dosimetry effect on total iron uptake was furthermore ruled out using THP-1-derived macrophages (Suppl. Fig.2a).

**Figure 2:**
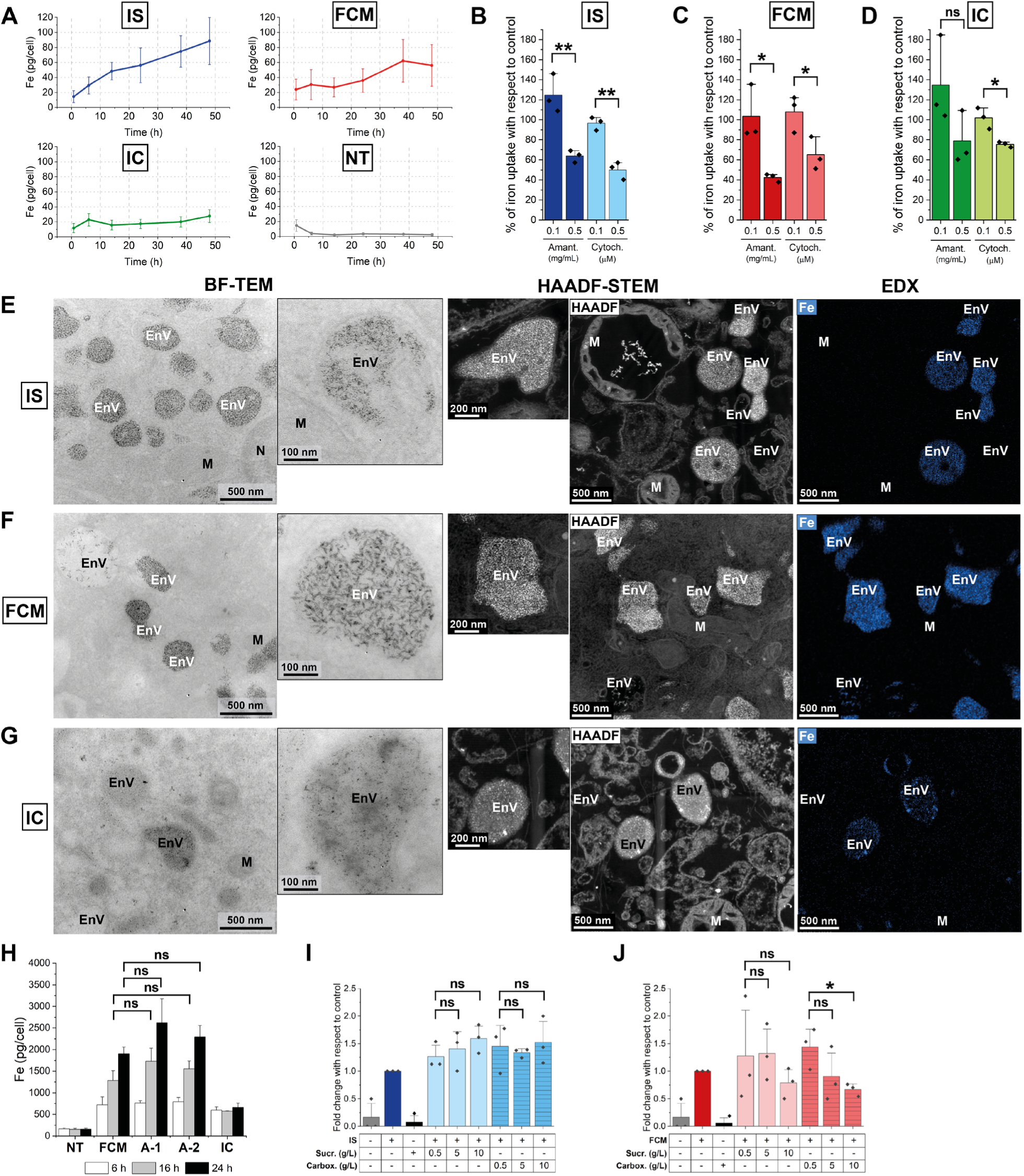
Dynamics of cell uptake, endocytosis and role of carbohydrate ligand. **A)** Total iron uptake by macrophages was quantified with ICP-OES over the indicated timepoints of treatment. The cells were exposed to 1800 µM of total iron of IS, FCM and IC. The data points represent the mean of the experiment performed in triplicates (with three different donors) ± S.E.M.. NT: non-treated. **B-D)** The endocytosis inhibitors Amantadine and Cytochalasin were used to pre-treat the cells for 60 min. Afterwards, **B)** IS, **C)** FCM or **D)** IC were added at a concentration of 1800 µM of total iron in the presence of the inhibitors. After 6 h of treatment, total iron was quantified with ICP-OES. The bar graphs represent the mean +S.E.M. of a triplicate with three different macrophage donors. *(0.05 > *P* > 0.01), **(0.01 > *P* > 0.001), ns: non-significant. **E-G)** Representative images of bright field (BF)-TEM (left) and scanning TEM (STEM) coupled with high-angle annular dark-field imaging (HAADF, middle) showing that **E)** IS and **F)** FCM are taken up by macrophages after 24 h into endocytic vesicles (EnV) as NPs with a morphology as seen in water and full medium (Fig.1, compare zoom-insets). These IS and FCM NPs have a clear iron signal (Fe) in elemental analysis *via* energy-dispersive X-ray spectroscopy (EDX, right). **G)** Amorphous, non-nanoparticulate precipitates of the Fe^3+^ IC control as in Fig.1 are also visible inside EnV. N: nucleus, M: mitochondria. Scale bars: BF-TEM: 500 nm, 100 nm; HAADF-STEM: 200 nm, 500 nm; EDX: 500 nm. **H)** The cells were treated for 6 h, 16 h and 24 h with FCM, A-1 (analogue 1), A-2 (analogue 2) and IC. Afterwards, total iron was quantified with ICP-OES. The bar graphs represent the mean + S.E.M. of a triplicate with three different macrophage donors. ns: non-significant. **I-J)** Competition assay between ICCs and their free carbohydrate ligands. The cells were pre-treated with either sucrose (Sucr.) or carboxymaltose (Carbox.) at the indicated concentrations. Then, **I)** IS or **J)** FCM were added for 6 h, and total iron content was measured with ICP-OES. The bar graphs represent the mean + S.E.M. of a triplicate with three different macrophage donors. The baseline iron content in the non-treated (NT) control cells stems from iron already present in the medium that was internalized by the cells. *(0.05 > *P* > 0.01), ns: non-significant.

Such different uptake dynamics likely reflect variations in the iron species within ICC formulations. Since NP-complexed fractions of IS and FCM are likely internalized *via* endocytosis, we used two endocytic inhibitors, Amantadine and Cytochalasin D, to evaluate endocytic activity in IS and FCM uptake (Fig.2b-d, Suppl. Fig.2b-e). While Amantadine is an inhibitor of clathrin-mediated endocytosis, Cytochalasin D depolymerizes the actin cytoskeleton. Both inhibitors reduced iron uptake of IS, FCM, and IC in a dose-dependent manner, with IC being least affected, indicating a distinct uptake mechanism and confirming its lower uptake rate (Fig.2a). The inhibitory effects of the Amantidine and Cytochalasin D were validated using an MTS assay and fluorescence beads (Suppl. Fig.2b). In addition, BF-TEM and scanning TEM (STEM) with high-angle annular dark-field imaging (HAADF) (Fig.2e-g, Suppl. Fig.2f-g) revealed IS and FCM NPs in intracellular endocytic vesicles (EnV), confirming macrophage uptake *via* endocytosis. The single spherical IS NPs measured 2-4 nm in length with few elongated clusters of 10-20 nm, and the ellipsoidal FCM clusters were of 10-25 nm length, aligning with their extracellular sizes (in water and in RPMI in Fig.1a-b, Table 1). Elemental analysis *via* energy-dispersive X-ray spectroscopy (EDX) confirmed the iron content in EnV of IS and FCM as NPs, while a less abundant, more localized iron signal was found in EnV for IC from its precipitates (Fig.2e-g, Suppl. Fig.2f-g).

To assess the contribution of LI to total iron uptake, we synthesized an analogue of FCM with double the LI content (A-1). After 6 h, 24 h and 48 h of treatment, no significant uptake difference was observed (Fig.2h), suggesting NP-complexed iron is the primary uptake determinant for FCM. A FCM analogue with double the particle size (A-2) showed no changes in uptake dynamics (Fig.2h).

In a previous study, proteomic and phospho-proteomic signatures of BMP2K ‒ a protein essential for clathrin-mediated endocytosis ‒ were differentially regulated in IS- and FCM-treated macrophages.^15^ This likely results from differences in composition and morphology of the carbohydrate ligand surface^7^ or other physicochemical properties affecting iron release. Since IS has a loosely bound sucrose ligand, while FCM features a strongly bound carboxymaltose ligand,^7^ we performed a competition assay between the ICCs and both of their free carbohydrate ligands. While sucrose did not interfere with the uptake of IS or FCM (Fig.2i), a pre-treatment with carboxymaltose decreased the total FCM uptake, but did not have a considerable impact on IS uptake (Fig.2j). These findings indicate that the interaction between IS and the cells is not mediated by the sucrose ligand, but likely *via* proteins forming the agglomerates we previously described (Fig.1). The fact that carboxymaltose partially blocked FCM uptake is in turn an indication that the FCM-cell interaction is indeed mediated by its carboxymaltose ligand, likely by binding to a membrane receptor with a carbohydrate-binding site. The inhibitory effect of carboxymaltose was furthermore specific to FCM, with no impact on other NPs (fluorescent beads, silica NPs, Suppl. Fig.2c-e).

In summary, IS and FCM exhibit distinct cell uptake profiles, driven by differences in their physicochemical properties. IS uptake likely involves multiple pathways: labile iron passively crossing the plasma membrane, and different endocytic pathways, depending on the iron species, e.g. TBI internalized *via* transferrin receptor (TfR)-mediated clathrin endocytosis and NP-complexed iron *via* bulk endocytosis ‒ a rapid process triggered by strong stimulation.^16,17^ In contrast, FCM, which consists primarily of NP-complexed iron, is predominantly internalized *via* bulk endocytosis. Notably, a complete endocytic event (e.g. clathrin-mediated) can occur within minutes.^18^ Hence, the delayed uptake observed for FCM with minimal internalization within the first 24 hours (Fig. 2a), suggests a considerable bottleneck in this step. We hypothesize that FCM’s initial interaction with its cellular receptor may involve low-affinity binding (as indicated by our UV-VIS data in Fig.1d) or require a co-receptor to facilitate uptake, thereby slowing down its internalization. Further studies are needed to confirm these mechanisms and identify potential factors influencing FCM’s uptake dynamics.

### *In vitro* PK profiling reveals a slower integration of FCM into the iron metabolism than IS due to a slower lysosomal digestion, controlling its bioavailability

Once inside the macrophages, the ICCs undergo a biodegradation process that has not yet been well characterized. After ferric iron (Fe^3+^) is mobilized from the ICCs, it is reduced to ferrous iron (Fe^2+^) joining the labile iron pool (LIP), followed by storage within tissue ferritin as Fe^3+^. The individual patient’s iron needs and homeostasis then drive signaling to export iron from ferritin to serum transferrin for transport to the bone marrow for red blood cell production.^6^

To characterize these steps, we conducted *in vitro* PK profiling using clinically relevant timepoints and readouts.^5^ To measure intracellular Fe^3+^ and Fe^2+^ content, we applied Prussian and Turnbull’s blue staining (with DAB enhancement) to the treated macrophages (Fig.3a). Comparing Fe^3+^ and Fe^2+^ levels based on these stainings over time revealed formulation-dependent biodegradation rates of ICCs (conversion from NP-bound Fe^3+^ iron to labile Fe^2+^ in endolysosomes, Fig.3b). Specifically, IS exposure showed a steady, parallel increase for both ions peaking at 38 h, while FCM exhibited delayed Fe^3+^ uptake and then still slower Fe^2+^ release than IS, indicating a postponed intracellular biodegradation of FCM independent of the uptake rate. These data are in line with previous findings that IS enables faster Fe²⁺ availability in cells compared to FCM.^19^

**Figure 3:**
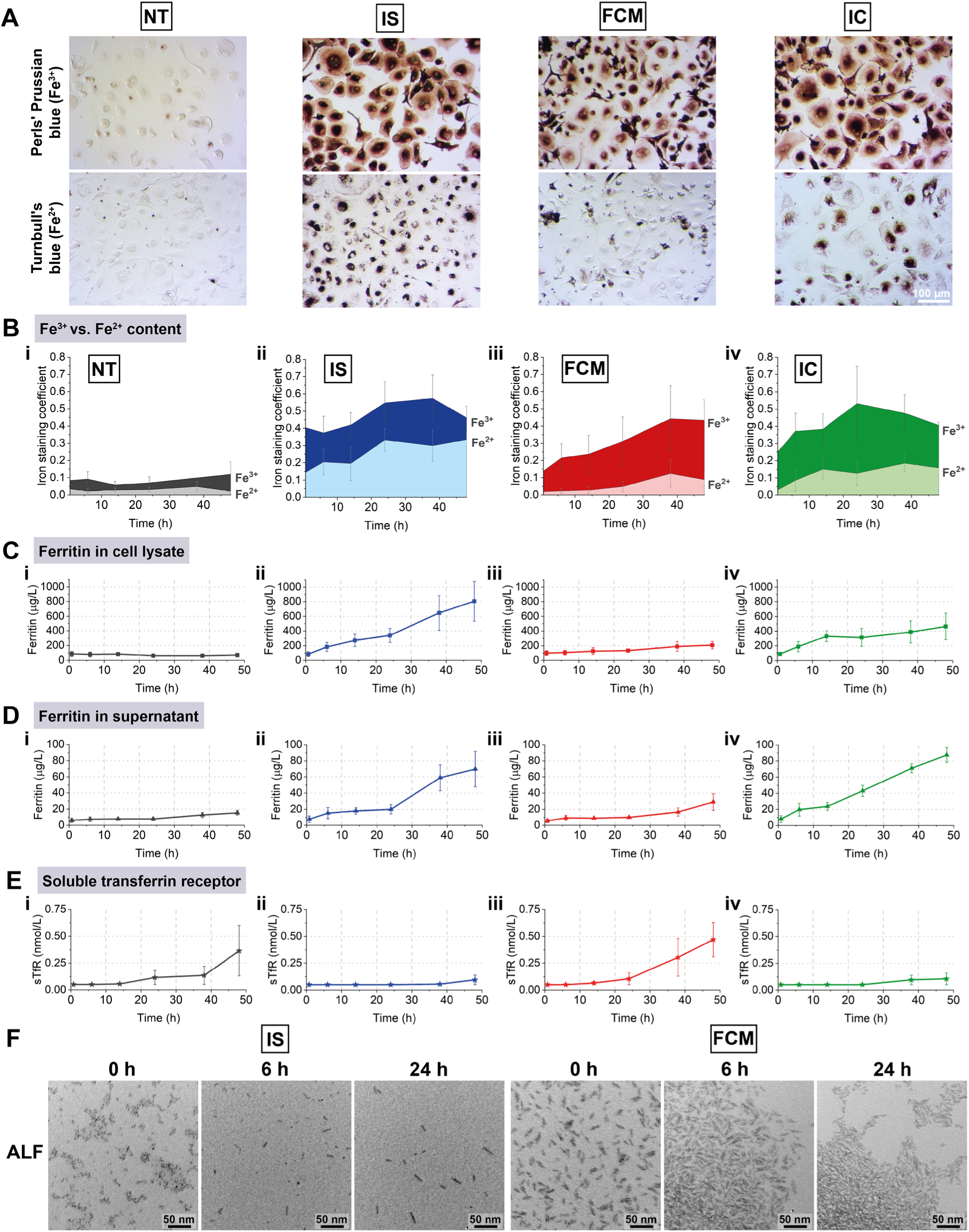
*In vitro* pharmacokinetic profiling of ICCs. Primary human macrophages were treated over time (0.75 h, 6 h, 14 h, 24 h, 38 h, 48 h) with IS, FCM or IC with 1800 µM of total iron. **A)** Representative light microscopy images of non-treated (NT) and IS/FCM/IC-treated (48 h) primary macrophages with Perls’ Prussian blue staining of Fe^3+^ and Turnbull’s blue staining of Fe^2+^, enhanced with DAB. Scale bar: 100 µm. **B)** Quantitative analysis of Fe²⁺/Fe³⁺ staining after cell fixation, staining and imaging with a brightfield microscope at each timepoint. The iron staining coefficient corresponds to the brown stained area normalized over the total cell area. The experiment was done in triplicates with cells from three different donors. **C-D)** Total human ferritin was quantified from the cell lysate and the supernatant. Additionally, soluble transferrin receptor (sTfR) released from the cells was measured in the supernatant **(E)**. All experiments were performed in triplicates with three different donors. The data points represent the mean ± S.E.M.. **F)** Representative BF-TEM images of IS and FCM NPs incubated over time in artificial lysosomal fluid (ALF, pH 4.5) showing that only elongated clusters of IS NPs are left after 24 h rotating at 37 °C, while the individual single spherical NPs are disappearing *via* dissolution to iron ions invisible with BF-TEM. Ellipsoidal FCM clusters instead retain their shape, size and abundance in endolysosome-mimicking ALF over time, showing even smaller inter-cluster distances, likely due to charge screening of the ligand charges by the salt ions of the ALF. Scale bars: 50 nm.

We confirmed this delay by quantifying ferritin and soluble transferrin receptor (sTfR) levels (Fig.3c-e), which are standard diagnostic indicators of the iron status in patients. Ferritin, an iron storage protein, increases with higher intracellular iron, while sTfR levels rise with iron deficiency.^20^ Following FCM treatment, macrophages began slightly producing ferritin at 24 h (Fig.3c-iii) and releasing it at 38 h (Fig.3d-iii), with a clear sTfR production at 38 h (Fig.3e-iii). IS, however, triggered ferritin production and release within 6 h (Fig.3c-d-ii), with no significant sTfR production (Fig.3e-ii), mirroring the Fe^3+^ control IC (Fig.3c-e-iv). These results indicate that FCM is internalized within the first 48 h (Fig.2a, 3a) but not immediately recognized as bioavailable iron, likely due to slowed biodegradation in endolysosomes. Indeed, BF-TEM imaging confirmed FCM’s stability over 24 h in artificial lysosomal fluid (ALF),^21^ which has a low pH of ∼ 4.5 and a high salt concentration, mimicking endolysosomal conditions (Fig.3f, Suppl. Fig.3). For IS, on the other hand, a disappearance of NPs was visible in ALF over 24 h. The stable carbohydrate moiety of FCM might more strongly resist the low pH and high salt concentration of the ALF compared to the more weakly bound carbohydrate ligand of IS, which could at least partially explain our observations, yet also other PCCs could influence FCM’s slower biodegradation rate.

### IS follows a standard and fast intracellular trafficking route with indications of transient cytotoxicity, while FCM accumulates in endosomes, delaying its biodegradation (*The Hamster Effect*)

To gain deeper insight into the ICC’s pathway from cellular uptake to endolysosomal biodegradation, we further analyzed the intracellular trafficking of ICCs (Fig.4, Suppl. Fig.4a). BF-TEM imaging revealed numerous densely packed EnV filled with IS NPs already at 45 min, in line with ICP-OES uptake measurements (Fig.1e). Over time, both the number and size of IS-filled EnV increased markedly (Fig.4a), and NP-loaded multi-vesicular bodies (MVBs) were particularly evident at 6 h and 24 h. Notably, swollen mitochondria were observed at early exposure time points (Fig.4c, Suppl. Fig.4b), suggesting oxidative stress in macrophages,^22^ likely triggered by rapid IS uptake, metabolization, and intracellular iron overload ‒ consistent with findings from our previous study.^15^ This mild toxic effect appeared to be reversible over time (48 h) and was specific to IS.

**Figure 4:**
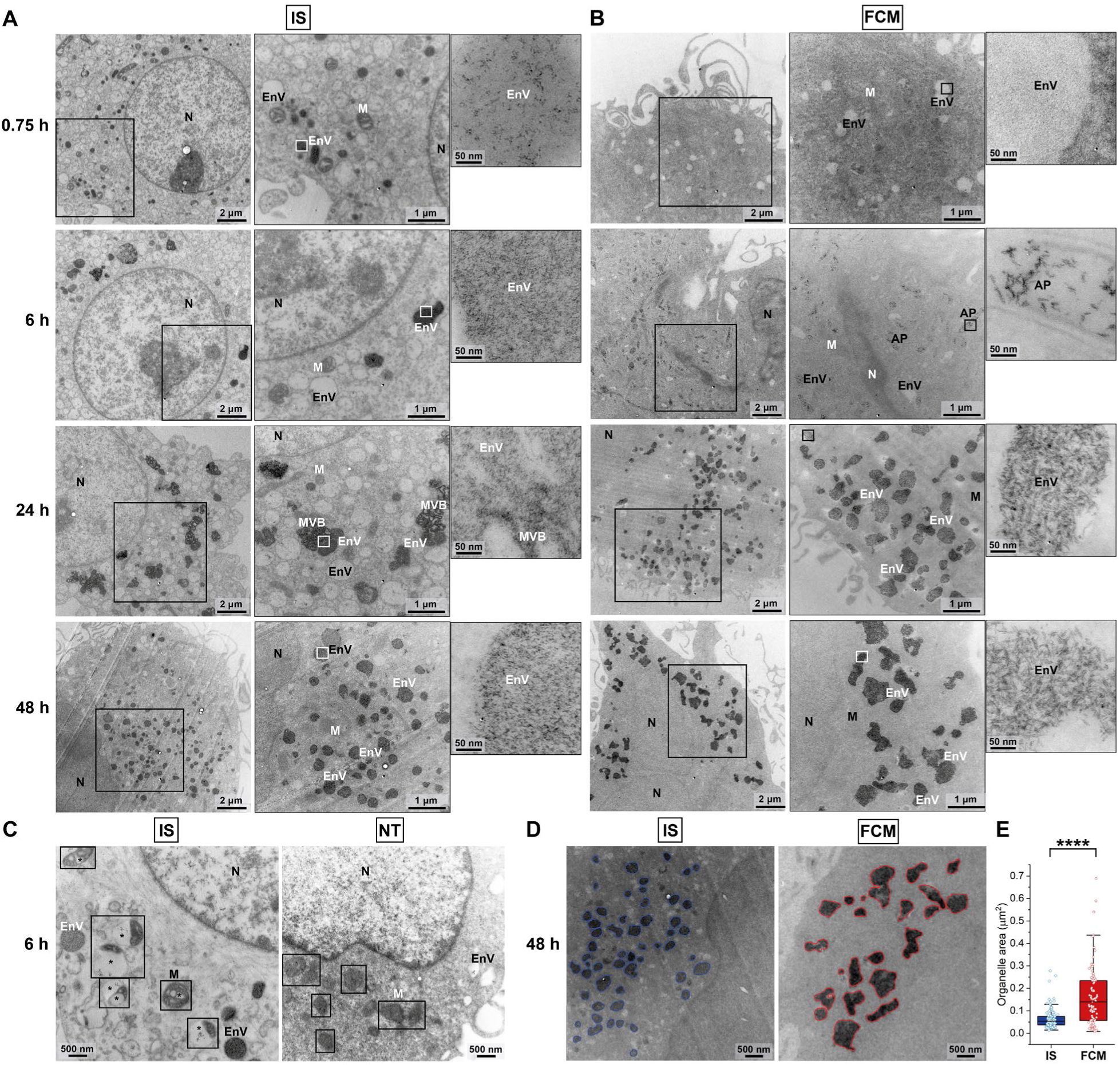
Intracellular trafficking. A-B) Representative BF-TEM images showing the intracellular trafficking of **A)** IS and **B)** FCM NPs in primary macrophages over time after uptake at different magnifications (black squares). **A)** IS NPs were internalized rapidly into endocytic vesicles (EnV), filling more EnV over time, and multi-vesicular bodies (MVB) loaded with IS NPs were observed. **B)** FCM NPs were taken up much later in time, sparsely filling EnV and autophagosomes (AP) at the beginning, and then forming densely packed, large, flower-shaped fused EnV until 48 h. Zoom-insets show that the morphology of IS and FCM NPs in EnV, MVB and AP is similar as in water and full medium (Fig.1). N: nucleus, M: mitochondria. Scale bars: 2 µm, 1 µm, 50 nm. **C)** At early time points, IS NPs led to swollen mitochondria (M, black squares) with obvious cavities between cristae (stars) visible in BF-TEM, indicative of oxidative stress, which was reversible over time and not observed to such extent in non-treated cells (NT). Scale bars: 500 nm. **D-E)** Instance segmentation of clearly visible, NP-filled EnV organelles as contoured in representative BF-TEM images **(D)** for IS and FCM after 48 h with EnV area quantification **(E)** indicating significantly larger FCM NP-filled EnV than with IS NPs. Scale bars: 500 nm. Mean ± S.E.M., ****P<0.0001.

In contrast, EnV loaded with FCM NPs were only detected from the 6 h time point onward (Fig.4b), aligning with the slower uptake rate of FCM (Fig.2a). At 6 h, a few NP-containing autophagosomes (AP), identifiable by their characteristic double-layer membranes, were observed (Fig.4b). Notably, by 24 h ‒ and even more prominently at 48 h ‒ large EnV, densely packed with FCM NPs, were evident (right panels in Fig.4b). These FCM-loaded EnV differed significantly in shape and size from their IS counterparts, displaying a distinctive flower-like structure and a much larger size (Fig.4d-e).

In the IC control, darker electron-dense precipitates were observed in EnV (black arrows), as also seen for IC in water and cell culture medium (Fig. 1a-b), which were absent in the untreated control (Suppl. Fig.4a). For both IS and FCM, the internalized and still visible NPs within EnV maintained their morphology and size over time, closely resembling their morphology in BF-TEM in water and medium (Fig.4a-b, zoom-insets; Fig.1a-b).

To further elucidate the intracellular fate of IS and FCM, and to identify which EnV are endolysosomes with active NP biodegradation, we developed a novel multi-modal approach combining BF-TEM imaging, elemental analysis *via* STEM-HAADF-EDX and Turnbull’s staining. Thereby, the Turnbull’s blue, which forms a dark precipitate in the presence of Fe^2+^ ions allowing for clear visualization with BF-TEM, served as a marker to identify compartments where NP biodegradation (and reduction to Fe^2+^) occurred (Fig.5). Accordingly, we classified the visible Fe^2+^-positive vesicles as endolysosomes (EL), i.e. where Fe^3+^ of the NPs is degraded to Fe^2+^, and Fe^2+^-negative vesicles as endosomes (E), which lack the low pH, and the high concentration of salts and enzymes required for iron biodegradation.

**Figure 5:**
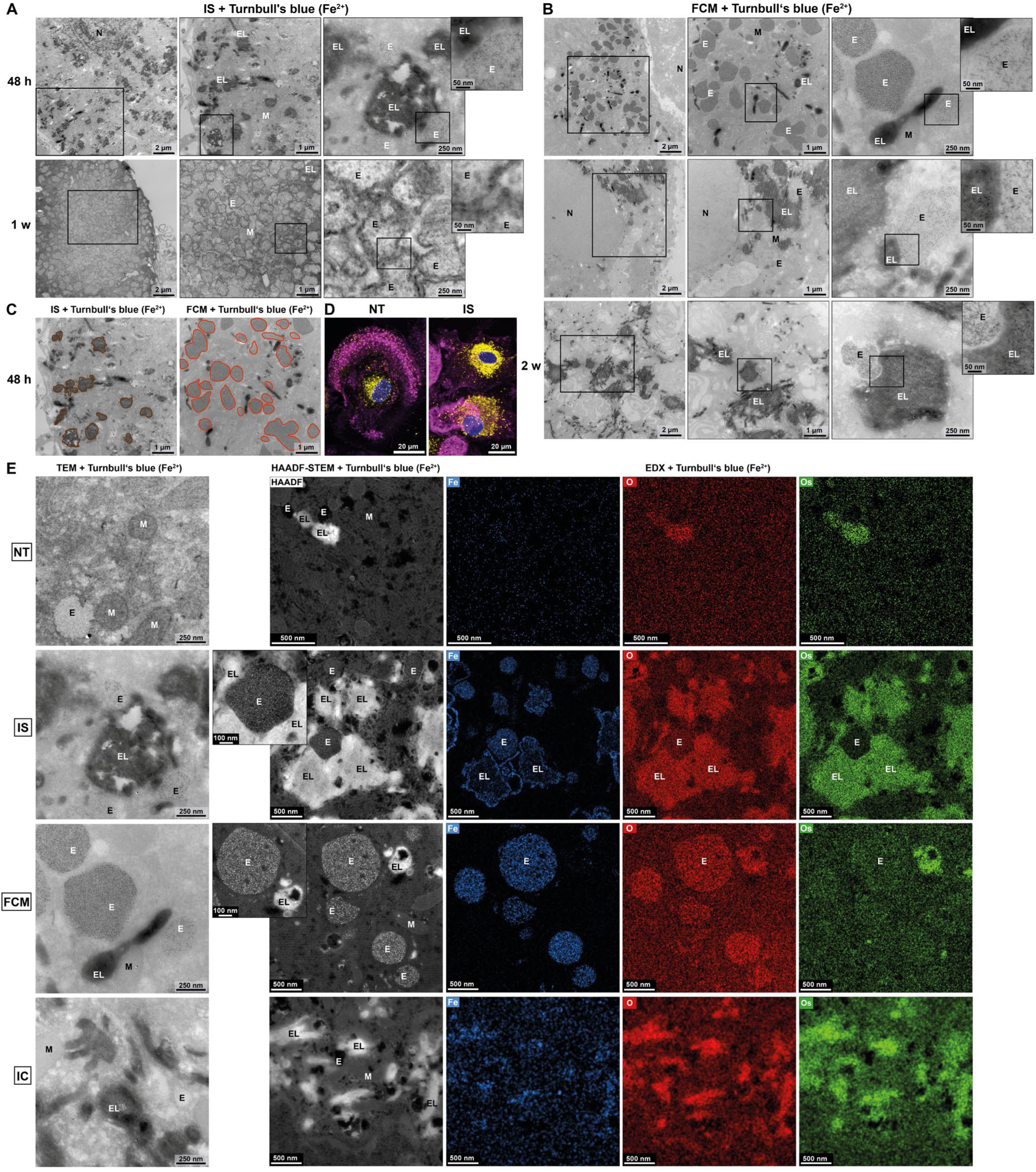
The Hamster Effect. **A-B)** Representative images of BF-TEM combined with Turnbull’s blue staining enhanced with DAB to visualize the digestion of Fe^3+^ in **A)** IS and **B)** FCM NPs to Fe^2+^ in endolysosomes (EL) at 48 h, 1 week (1 w) and 2 weeks (2 w) after uptake by primary macrophages at different magnifications (black squares). **A)** For IS NPs, many large and strongly Turnbull’s-stained EL (dark structures) were visible at 48 h as well as much smaller IS NP-filled endosomes (E) without iron digestion and thus Turnbull’s staining. After 1 week, the cells showed many IS NP-filled EnV with ruptured membranes, indicative of beginning cell death, and dark Turnbull’s staining of the cytoplasm. The IS-NP treated cells did not survive until 2 weeks. **B)** For FCM NP-treated primary macrophages, the large and strongly Turnbull’s-stained EL (dark structures) as seen for IS NPs at 48 h were only visible at 1-2 weeks and still contained more undigested FCM NPs (compare zoom insets). FCM NPs remained longer in E with delayed fusion to EL and iron digestion, leading to densely packed, flower-shaped fused E at 48 h that diminished in size until 2 weeks but kept their irregular shape (see 2 w). The IS and FCM NPs showed a similar morphology as in water or medium (Fig.1) where visible in E and EL (zoom insets). N: nucleus, M: mitochondria. Scale bars: 2 µm, 1 µm, 250 nm, 50 nm. **C)** Contoured EL with IS NP-digestion to Turnbull’s-positive Fe^2+^ and large FCM NP-filled, flower-shaped E without digestion in BF-TEM combined with Turnbull’s blue staining highlight the size difference of the EnV and the absence of Turnbull’s-stained EL for FCM NP-treated primary macrophages at 48 h. Scale bars: 1 µm. **D)** Representative immunofluorescence images of non-treated (NT) macrophages or treated with IS for 48 h. Staining: Lamp1 (yellow), actin (magenta), nuclei (blue). Scale bars: 20 µm. **E)** Combination of representative BF-TEM images (left) as in **A-B)** with Turnbull’s staining (enhanced with DAB) and HAADF-STEM-EDX (middle, right) at 48 h revealing that the Turnbull’s-stained and thus Fe^2+^-positive EnV are EL with a characteristic osmium signal (Os) from the high concentration of endosomal digestive enzymes, and with iron (Fe) and oxygen (O) signals from still undigested IS/FCM NPs inside the EL. FCM NP-filled EnV are devoid of osmium signal, indicating that they are E without digestive enzymes. Upon IC-treatment, localized Fe and O signals of IC precipitates in EL with Os signal are visible, while in non-treated cells the Fe background signal is diffusely distributed over the entire cell, with O and Os signals localizing to EL as expected from their high digestive enzyme and salt concentrations. Scale bars: 250 nm (BF-TEM); 100 nm, 500 nm (HAADF-STEM-EDX).

For IS, numerous Fe^2+^-positive endolysosomes were clearly observed at 48 h, often in close proximity to fewer and much smaller Turnbull’s-negative endosomes filled with IS nanoparticles (Fig.5a,c). Lamp1 immunofluorescence imaging of IS-treated macrophages confirmed an increased number of endolysosomal vesicles (yellow) compared to the non-treated control (Fig.5d), supporting the active degradation of IS NPs within endolysosomes.

HAADF-STEM-EDX further confirmed the identity of these compartments (Fig.5e, Suppl. Fig.5c), revealing distinct elemental profiles: Fe^2+^-positive, Turnbull’s stained endolysosomes displayed strong iron and oxygen signals from the yet undegraded ICC NPs, as well as high osmium (and nitrogen) signals indicating the high concentration of endolysosomal digestion enzymes, whereas NP-filled endosomes without such enzymes exhibited the ICC’s iron and oxygen signals but low osmium (or nitrogen) levels.

Interestingly, the flower-shaped, FCM-loaded vesicles were negative for Turnbull’s staining, and exhibited high iron and oxygen content with low osmium (or nitrogen) signals ‒ characteristic of endosomes rather than endolysosomes (Fig.5b,c,e). This suggests that FCM NPs accumulate in endosomes for an extended time prior to endosomal fusion with lysosomes. We termed this phenomenon the *Hamster Effect*, drawing an analogy to how hamsters store food in their cheek pouches. Similarly, macrophages internalize FCM NPs into endosomes but, instead of fusion with lysosomes and immediate digestion of FCM NPs in endolysosomes, they temporarily sequester FCM NPs in enlarged endosomes. This storage phase delays FCM NP biodegradation, leading to a gradual and sustained release of iron over time. As expected, IC exposure resulted in Fe^2+^-and Turnbull’s-positive endolysosomes that lacked nanoparticles (Suppl. Fig.5a).

The *Hamster Effect* persists for approximately one week: large Fe^2+^-positive endolysosomes still containing undegraded FCM NPs began to appear after one week of exposure and increased in abundance over two weeks, indicating a transition from enlarged endosomes (visible at 48 h) to Turnbull’s stained FCM-digesting endolysosomes (Fig.5b). Supporting this shift, FCM-filled endosomes without Turnbull’s staining gradually decreased in size during this period, while retaining their flower-shape. IS-treated macrophages exhibited ruptured EnV membranes after 1 week and cell death after two weeks. In contrast, FCM-treated cells remained viable for longer (up to 2 weeks), suggesting that while IS uptake and digestion led to rapid iron overload and cytotoxicity, FCM was processed gradually, ensuring a sustained and more controlled iron bioavailability.

The slower intracellular processing of FCM compared to IS is consistent with observations for other nanoparticle systems, such as GalNAc–conjugated siRNAs^23^ and 30 nm polystyrene beads^24^. The latter have been shown to form large intracellular vesicles in macrophages, preventing transport from endosomes to endolysosomes.^24^ In the case of FCM, this delayed intracellular trafficking ‒ referred to as the *Hamster Effect* ‒ can be attributed to the formation of large flower-like vesicles seen in TEM, where FCM nanoparticles remain sequestered for extended periods without degradation. To further investigate the mechanisms underlying the *Hamster Effect*, we analyzed the expression of early (EEA1) and late (Rab7) endosomal markers by Western blot following 6- and 48-hour treatments with IS and FCM (Fig.6a-e). While FCM induced an increase in Rab7 at 6 hours (Fig.6e), this effect was highly donor-dependent, and no significant differences were observed in overall protein expression levels. Since the total expression levels of early and late endosomal proteins seemed unchanged, we next explored the morphology and density of intracellular vesicles stained with the early endosomal marker EEA1, the late endosomal protein Rab7 or the endolysosomal marker Lamp1 in immunofluorescence (Fig.6f-m), which allows the visualization of endocytic vesicles at different stages. Primary human macrophages exhibit considerable morphological heterogeneity (Fig.6f), yet distinct intracellular vesicles expressing EEA1, Rab7, or Lamp1 were detected following IS and FCM treatments over 48 h (Fig.6g). Both IS and FCM led to a significant increase in the number of vesicles carrying the early endosomal marker EEA1 and the late endosomal protein Rab7 from 6 to 48 h compared to untreated cells (****P_FCM, no. objects_, **P_IS, no. objects_, Fig.6h,k; ***P_IS,FCM, no. objects_, Fig.6i,l). IS further showed enlarged Rab7- and Lamp1-positive vesicles, i.e. late endosomes and endolysosomes, compared to untreated cells within 48 h (***P_IS, total area_, ***P_IS, avg. object area_, Fig.6i; ^ns^P_FCM, avg. object area_, Fig.6j), while FCM caused a moderate Rab7 vesicle enlargement (***P_FCM, total area_, ^ns^P_FCM, avg. object area_, Fig.6l) and only a slight increase in Lamp1 vesicle size (^ns^P_FCM, avg. object area_, Fig.6m). These findings suggest that IS triggers the maturation of early to late endosomes, their fusion to endolysosomes and their enlargement, while the flower-like *Hamster* vesicles observed for FCM within 48 hours still possess more of an endosomal rather than an endolysosomal identity, aligning with our EDX data (Fig.5).

**Figure 6:**
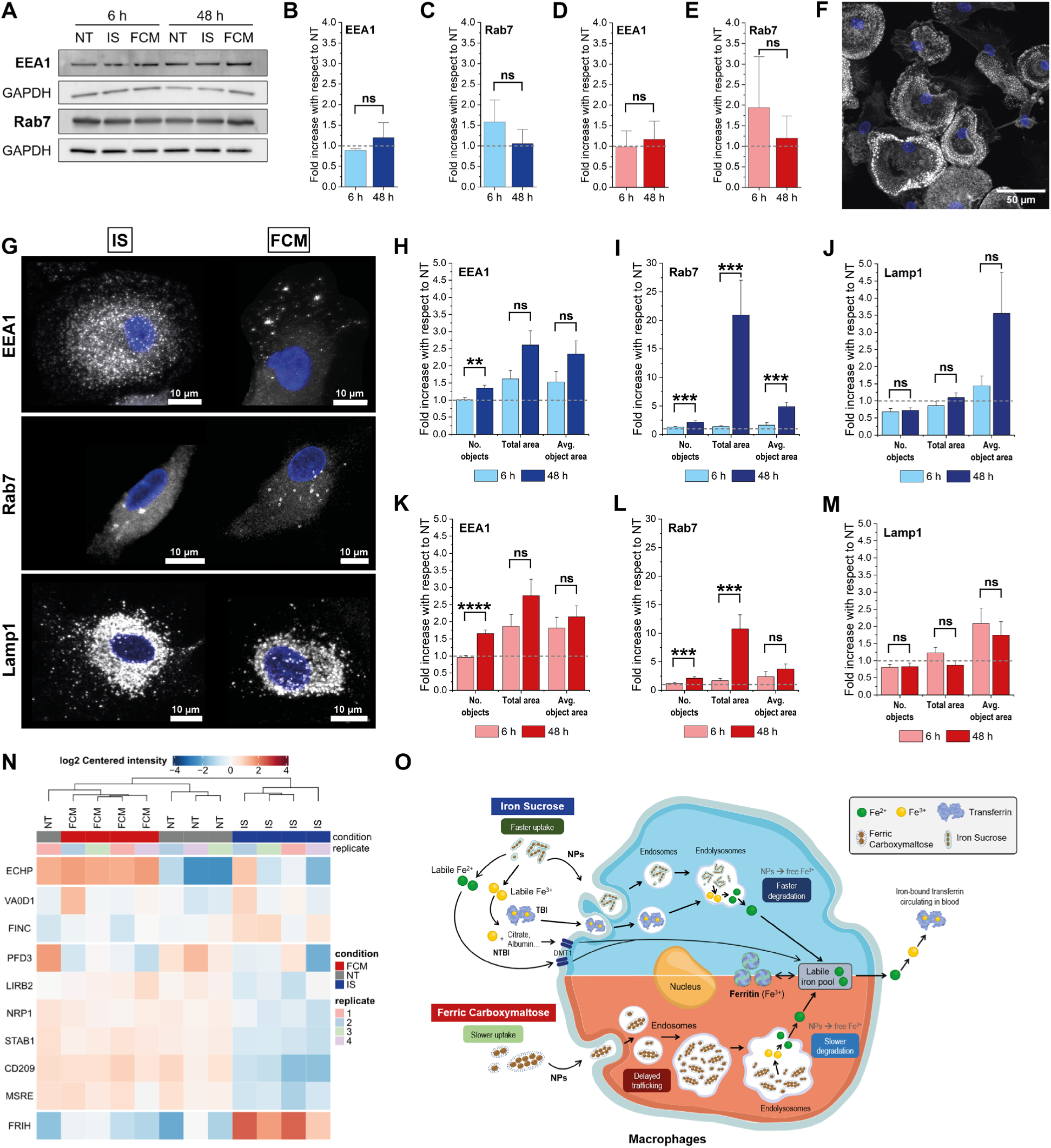
Mechanism of action of ICCs in macrophages. A-E) The total EEA1 and Rab7 protein expression was quantified by Western blot. **A)** Cropped blots of one donor are shown with GAPDH as loading control. See Supp. Fig. 6a-b for the complete blots. **B-E)** Quantification of the Western blots as in **A)** conducted in triplicates (with three different donors). The bar graphs represent the mean + S.E.M.. ns: non-significant. **F-M)** The morphology and density of intracellular vesicles was analyzed using confocal fluorescence microscopy. **F)** Actin staining (phalloidin; gray) shows the heterogeneous morphology of the cells. Nuclei are shown in blue. Scale bar: 50 µm. **G)** Images of macrophages treated with IS or FCM for 48 h, and stained for EEA1, Rab7, or Lamp1 proteins (gray). Nuclei are shown in blue. The images are a maximum projection generated with ImageJ. Scale bars: 10 µm. **H-M)** The size and density of vesicles expressing either EEA1, Rab7 and Lamp1 was quantified from the images as in **F-G)**. *No. objects* represents the number of stained objects (contours) identified in the image. *Total stained area (Total area)* refers to the overall area covered by the detected objects. *The average object area (Avg. object area)* represents the average size of the detected objects. The bar graphs show the mean + S.E.M. of 60 to 110 cells. The data was normalized to the non-treated control of its corresponding donor. The experiment was done in triplicates with three different macrophage donors. *(0.05 > P > 0.01), **(0.01 > P > 0.001), ns: non-significant. **N)** Proteomics heatmap showing the log2 centered intensity of the proteins detected as significantly differentially expressed in at least one of the comparisons after 6 h of IS or FCM treatment *versus* untreated primary human macrophages. **O)** We propose a novel hypothesis for ICC processing in macrophages. Iron sucrose (IS) nanoparticles are rapidly taken up and degraded in endolysosomes, causing a quick rise in bioavailable iron but also oxidative stress. In contrast, ferric carboxymaltose (FCM) follows a slower route with delayed uptake and degradation, leading to gradual iron release ‒ termed the *Hamster Effect* ‒which may explain its lower toxicity. In both cases, the iron is eventually stored in ferritin, enters the blood, binds to transferrin, and is transported to the bone marrow. LSEC: liver sinusoidal endothelial cells. DMT1: divalent metal transporter 1.

Next, we addressed the question how FCM can induce the formation of such enlarged, morphologically altered *Hamster* vesicles without significantly increasing the endosomal protein expression (EEA1 and Rab7; non-significant in Western blot, Fig.6d-e; non-significant total area and average object area changes in immunofluorescence, Fig.6k,l). Proteomics analysis identified Enoyl-CoA Hydratase/Isomerase (ECHP) as a key differentially expressed protein following a 6-hour FCM treatment (Fig.6n). ECHP catalyzes the hydration and isomerization of enoyl-CoA intermediates during fatty acid β-oxidation, generating acetyl-CoA ‒ a precursor for lipid biosynthesis.^25^ This could indicate an increased lipid biosynthesis in FCM-treated macrophages, which in turn may enable the formation of the enlarged endosomal vesicles detected in TEM at 48 hours that do not have EEA1 or Rab7 protein levels proportional to their vesicle size (Fig.4-6). Notably, FCM-treated macrophages also exhibited an upregulation of transmembrane receptors containing C-type lectin-like domains, including NRP1, STAB1 and CD209, along with the macrophage scavenger receptor MSRE. These findings suggest that these multi-functional cell-surface receptors may eventually be repurposed by macrophages for FCM nanoparticle endocytosis *via* carbohydrate ligand-receptor interactions, supporting our observation that FCM is not immediately recognized by the cells due to the lack of canonical, high affinity-binding receptors for FCM. Furthermore, fibronectin (FINC) is rather downregulated in FCM-treated macrophages, potentially lowering the binding of opsonins to FCM NPs and thereby further hampering their recognition for uptake and degradation, in contrast to IS NPs. Interestingly, IS strongly upregulated the expression of the ferritin heavy chain protein (FRIH), indicating a stronger demand for iron storage as ferritin in IS-treated macrophages at 6 hours, consistent with our other assays.

While more in-depth studies are needed to further elucidate these underlying mechanisms, our results provide unprecedented insights into the metabolic fate of ICCs depending on their physicochemical properties and deliver likely explanations for their different pharmacokinetic and pharmacodynamic profiles observed *in vivo*.

## Conclusions

This study provides a mechanistic understanding of how different ICCs interact with macrophages, revealing key insights into their intracellular fate. Using a combination of advanced microscopy, elemental analysis, and *in vitro* pharmacokinetic profiling, we demonstrate that ICC uptake, intracellular trafficking, and biodegradation are highly dependent on their physicochemical characteristics.

We propose a novel hypothesis for ICC processing in macrophages (Fig.6o). IS undergoes rapid endocytosis as nanoparticles (NPs) but also as other iron species (TBI – Transferrin-bound iron, NTBI – Non-transferrin-bound iron, labile Fe^2+^). Uptake of IS NPs is followed by early endolysosomal trafficking, where it is quickly degraded, leading to a sharp increase in bioavailable iron. The rapid metabolism of IS, however, can induce oxidative stress, as evidenced by swollen mitochondria at early and membrane rupture at later time points. In contrast, FCM follows a distinct intracellular route, characterized by delayed uptake and sequestration to enlarged endosomes, a phenomenon we term the *Hamster Effect*. Macrophages internalize FCM NPs, but postpone their degradation, delaying iron mobilization. This process results in a more gradual and sustained iron release, explaining FCM’s slower pharmacokinetics and lower cytotoxicity compared to IS. For both IS and FCM, bioavailable iron is incorporated into the labile iron pool (LIP) and stored associated to ferritin. Finally, it is released from the macrophages and enters the blood circulation, where it will bind to transferrin and is transported to the bone marrow. This process is aligned with FCM’s better stability profile and slower clearance from plasma *in vivo*, allowing the administration of larger single doses of FCM compared to IS.

Our findings provide the first direct evidence of distinct ICC processing profiles in macrophages, challenging the assumption that all ICCs follow the same biodegradation pathway. This mechanistic insight now begins to bridge the gap between the physicochemical properties of nanoparticles and their pharmacokinetic profiles, offering a framework for optimizing ICC treatment in patients with iron deficiency anemia.

## Acknowledgements

We thank the Scientific Center for Optical and Electron Microscopy (ScopeM) of ETH Zurich, specifically Stephan Handschin for providing support and access to their ultramicrotomes, and to Christian Zaubitzer for access to the Talos 200 electron microscope and performing few EDX-STEM-HAADF measurements for this study.

## Funding

This study was co-funded by the Empa, St. Gallen, Switzerland and CSL Vifor, Switzerland. IS and FCM nanoparticles were an in-kind gift from CSL Vifor.

### Author contributions

Conceptualization: V.A.N., P.W., R.D., A.E.A., B.F. Investigation, methodology, validation: S.E., V.M.K., A.R., L.K., J.J.B., S.G., A.G., D.B., K.H.S., V.A.N. Data curation: S.E., V.M.K., A.R., V.A.N. Formal analysis: S.E., V.M.K., A.R., L.K., D.B., K.H.S., J.Bo., R.R.P., V.A.N. Software: D.B., K.H.S. Project administration: V.A.N. Supervision: V.M.K., V.A.N., and P.W. Funding acquisition, resources: P.W., B.F. Visualization: V.M.K., V.A.N. Writing—original draft: S.E., V.M.K., V.A.N. Writing—review & editing: S.E., V.M.K, A.R., R.D., A.E.A., B.F., P.W., V.A.N.

## Methods

### 1. Cell Culture

Human primary macrophages were isolated from buffy coats of healthy adult donors obtained from the blood bank in Zurich, Switzerland (ethical approval BASEC Nr. Req_2021-00687). First, peripheral blood mononuclear cells (PBMCs) were extracted from these buffy coats using the Ficoll (Sigma-Aldrich) density gradient centrifugation protocol. After negative selection of PBMCs with a Classical Monocyte Isolation Kit (Miltenyi Biotec), monocytes were differentiated into M0 macrophages in RPMI-1640 (Sigma-Aldrich) culture medium (supplemented with 10% fetal calf serum (FCS), 1% penicillin-streptomycin (PS)) and 20 ng/ml M-CSF (Gibco) for 7 days at 37 °C, followed by a M2 macrophage polarization period of 1.5 days in fresh cell culture medium containing IL-4 (20 ng/ml) and IL-13 (20 ng/ml) (Miltenyi Biotec).

THP-1 cells were seeded in RPMI-1640 culture medium (Sigma-Aldrich) supplemented with 10% fetal calf serum (FCS), 1% penicillin-streptomycin (PS), and 2 mM L-glutamine, in the presence of 200 nM PMA. After a 3-day incubation, the PMA-containing medium was removed, and cells were allowed to rest for 24 hours before being used in experiments.

### 2. Iron Formulations

Iron carbohydrate complex (ICC) treatments Iron sucrose (IS, Venofer®, Lot No. 1881022C, CSL Vifor) and Ferric Carboxymaltose (FCM, Ferinject®, Lot No. 0970022E, CSL Vifor), as well as Iron citrate (Sigma Aldrich, 2338-05-8) as a non-NP-complexed iron control, were used in our experiments. On the day of the experiment, these iron formulations were extracted from sealed vials using a syringe to prevent oxidation and diluted in RPMI-1640 with 10% FCS and 1% PS to achieve a total iron concentration of 1800 μM. This concentration mirrors the expected plasma iron levels following the administration of 300 mg of IS in patients and is therefore a clinically relevant iron concentration.

### 3. Perls’ Prussian blue, Turnbull’s blue and DAB staining

Perls’ Prussian blue staining^26^ was performed for cells under static and dynamic conditions. Perls’ Prussian Blue stain is widely utilized for the detection of iron (Fe^3+^) in fixed cells and tissues, while the Turnbull’s blue^27^ stain is used to visualize iron (Fe^2+^). All experiments were performed in triplicates to ensure reproducibility.

#### 3.1 Staining of Cells

After treatment with ICC, the cells were washed three times with PBS (Sigma, Germany) and fixed with 4% paraformaldehyde (PFA) for 20 minutes at room temperature (RT). The fixed cells were then exposed Prussian blue Perls’ reagent (4% potassium ferrocyanide 12% HCl, 1:1 v/v ratio) or Turnbull’s blue reagent (4% potassium ferricyanide/12% HCl, 1:1 v/v ratio) for 6 hours at room temperature, followed by three PBS washes. To further enhance the iron staining signal, 3,3’-diaminobenzidine (DAB, Sigma Aldrich) staining was used. For this, cells were treated with 0.05% DAB in PBS for 10 minutes at room temperature. Subsequently, the solution was replaced with 0.05% DAB in PBS with 0.033% H_2_O_2_ and incubated for an additional 10 minutes at room temperature. The cells were washed three times with PBS, imaged using a light microscope (Zeiss) with 5-10 images captured per condition for data representation, and then stored at 4 °C.

#### 3.2 Prussian Blue - Dynamic Conditions

In a black 96-well plate with a clear bottom (Greiner, Lot No. E24053E8), 100 µl iron formulations were prepared as follows: 500 µM of IS, FCM or Iron citrate at concentrations of 250, 500, 750, and 1000 µM and 0.14, 1.4, and 2.8 g/l of Sucrose or Carboxymaltose (free carbohydrate ligands of IS and FCM, respectively) with or without 500 µM IS or FCM. To initiate the staining, 100 µl of Prussian Blue stain (4% potassium ferrocyanide 12% HCl, 1:1 v/v ratio) was added per well. The reaction was monitored over 120 minutes at 37 °C by measuring absorbance at 650 nm at 53-second intervals in a plate reader (BioTek, Synergy H1).

### 4. Quantitative Image Analysis

#### 4.1 Fe^2+^/Fe^3+^ Staining

The quantitative image analysis of Fe²⁺/Fe³⁺ staining was performed in two steps in Matlab (R2023b): first, a staining detection step, followed by a cell area detection step. Staining detection was performed by applying a dynamic threshold (ROI ≤ xmax-50) on the brightfield images, with xmax representing the histogram maximum. This threshold proved to effectively exclude white background pixels and selectively include the stained area in the detected region of interest. The number of pixels in the detection was determined as the stained area. Confluency detection was performed by applying edge detection and a dilation step, followed by filling interior gaps and smoothing the objects in the detection. The objects in the detection were statistically analyzed, and outliers were excluded to remove artifacts and debris. Finally, the edges of detected objects were smoothed before determining the number of pixels in the detection as the total cell area. The staining coefficient was determined as the stained area normalized over the total cell area.

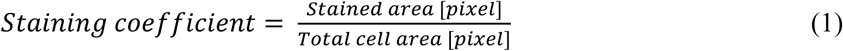

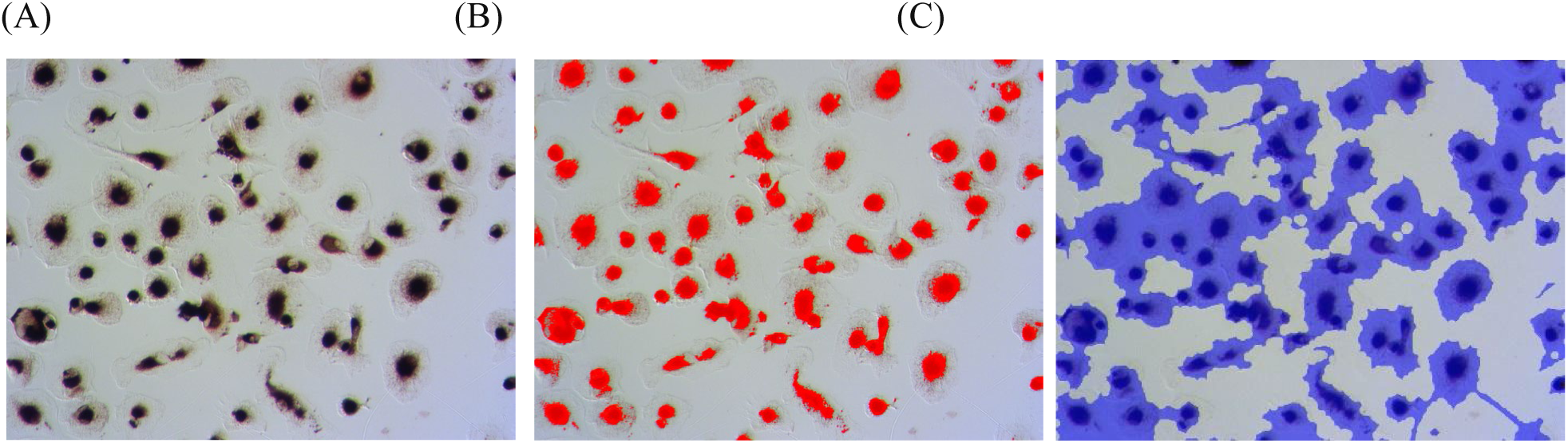

Fe^2+^/Fe^3+^ staining analysis. (A) Original image. (B) Staining detection. (C) Cell area detection.

#### 4.2 Endolysosomal Stainings

Fluorescent images were analyzed using Python v3.12.3. Colocalization of the different stainings was quantified using Pearson’s correlation and Mander’s overlap coefficient. For actin, EEA1, Lamp1 and RAB7 staining, positive signals were extracted using thresholding. Shape descriptors of the objects within the mask were extracted, as well as the location relative to the DAPI-stained nuclei. Cluster analysis of stained objects was performed using DBSCAN.

### 5. Inductively Coupled Plasma - Optical Emission spectroscopy (ICP-OES)

ICP-OES was used to quantify the total iron content within cells. Human primary monocytes were seeded in 6- or 12-well plates (TPP) at a density of 1 Mio cells/well, differentiated, polarized and treated as described above. Cells were washed three times with PBS and lysed for 20 minutes at 4°C in 700 µl Lysis buffer (150 mM NaCl, 1% Triton-X, 50 mM Tris-HCl, pH 7.4). 630 µl of this sample, 147 µl of 68% HNO_3_, and 170 µl of H_2_O_2_ were added to a Teflon tube, to be digested in a microwave suitable for elemental analysis to ensure complete sample breakdown. After digestion, the sample was brought to a volume of 5 ml by dilution with Milli-Q water and calibration standards were prepared in 2% HNO_3_ from 0.1 to 10 ppm (0, 0.1, 0.5, 1, 5, 10 ppm) by diluting the Fe stock ion solution (CGFE1, Inorganic Ventures, Lot. No. U2-FE734834). Measurements were performed using the axial measurement mode of a 5110 ICP-OES instrument (Agilent Technologies).

From the remaining 70 µl of lysed cells, total DNA was quantified by the Quant-iT™ PicoGreen™ dsDNA Assay Kit (ThermoFisher) following the manufacturers protocol. The DNA quantity, expressed in picograms (pg), served as a metric for determining the iron content per cell (pg Fe/cell) by calibrating the iron and DNA levels within the sample. To this end, it was approximated that a single cell comprises 6 pg of DNA.

### 6. Detection of Labile Iron in ICCs Diluted in Cell Culture Media

In prewarmed RPMI-1640 cell culture media or Milli-Q water, 1800 µM of iron formulations (IS/FCM/IC) were added and incubated for 0.75 or 24 hours at 37 °C. A minimum of 5 ml of treatments were filtered in 30 kDa, 15 ml Ultra-15 Centrifugal Filter Units (Millipore, UFC903024) by 30-minute centrifugation at 3000 RCF (MPW-380R, SN13/17 REF 11776 1400RPM). Filtered samples were collected and analyzed by ICP-OES.

### 7. Inhibition of Endocytosis

Polarized cells were pretreated for 1 hour with 0.1/0.5 mg/ml Amantadine (Sigma, A1260), 0.1/0.5 µM Cytochalasin D (Sigma, C8273), and 0.1% DMSO as a control, before they were treated for 6 hours with 1800 µM iron formulations (Iron Sucrose/Ferric Carboxymaltose) mixed with these inhibitors at 37°C. The supernatant was collected in a 15 ml Falcon tube, centrifuged at 300 g for 10 minutes to rescue any floating cells. Cells were washed three times with PBS, lysed and further processed following the ICP-OES protocol. All experiments were performed in triplicates to ensure reproducibility.

### 8. FCM Analogs

Two different FCM analogs were synthesized with different manufacturing processes leading to alterations in physicochemical characteristics. Analog 1 exhibits a higher hydrodynamic size and molecular weight compared to the FCM reference listed drug (RLD). Analog 2 exhibits a higher Fe (II) compared to FCM RLD. Polarized cells were treated for 6, 16 or 48 hours with 1800 µM iron formulations and two FCM analogs. Cells were washed, lysed, and processed per ICP-OES protocol. All experiments were performed in triplicates to ensure reproducibility.

### 9. Impact of ICC Sugar-Shell Ligands on Iron Formulation Uptake

To assess impacts on cellular iron formulation uptake following exposure to varying concentrations of sugar-shell ligand solutions, 0.5, 5 or 10 g/ml Sucrose or Carboxymaltose were diluted in RPMI-1640 cell culture media. After 30 minutes of pretreating M2a macrophages with this solution, 1800 μM iron formulations (Iron Sucrose/Ferric Carboxymaltose) were added and cells were incubated at 37 °C for 6 hours. After washing thrice with PBS, samples were processed further following the ICP-OES protocol.

### 10. Bright-Field Transmission Electron Microscopy (BF-TEM)

In a 12-well plate (TPP), 0.55 Mio human primary monocytes were seeded on 18 mm cover slips (VWR, HECH40990121), differentiated, polarized and treated with 1800 µM iron formulations for 0.75, 6, 24, and 48 h. For the long-term experiments cells were treated for 48 h with 1800 µM iron formulations, washed once with fresh media and then cultivated in media without iron formulations up to 2 weeks until fixation. During this time, media was exchanged twice a week. At the respective time points, the samples were washed in PBS trice and fixed in a 1% glutaraldehyde and 4% PFA solution for 40 minutes at room temperature. For samples stained with the Turnbull’s and DAB staining, the same protocol was used as described before in static conditions after this fixation step. After DAB staining, cells were washed twice with PBS and twice with 0.1 M Na-cacodylate buffer. Turnbull’s blue stained and unstained samples were contrasted with 1% osmium tetroxide (Electron Microscopy Sciences) in 0.1 M Na-cacodylate buffer for 1 hour at room temperature in the dark. Cells were then washed trice in Milli-Q water for 5 minutes at room temperature before being dehydrated with an ethanol gradient of 30% (5 min), 50% (5 min), 70% (5 min), 90% (5 min) and 100% (10 min, trice) and incubated in a 1:1 solution of 100% Ethanol and Epon for 1 hour at room temperature. After an incubation in fresh Epon over night at room temperature with open lid (for Ethanol evaporation), the coverslips were removed from the wells and mounted onto microscopy glass slides with a few drops of Epon (the cells facing up). Epon-filled gelatin capsules (Polysciences Inc, 07348-1000) were then placed on top, and the assembly was cured at 55-60 °C for approximately 72 hours. The hardened capsules with cells were removed from the glass slides on a heating plate at 60-70 °C and 60-80 nm thin sections were cut using an Ultramicrotome (Leica EM UC6) with an ultra 35° diamond knife (DiATOME). Sections were deposited on Formvar-coated copper grids (100 hexagonal mesh, EM Resolutions).

To visualize the iron formulations in MilliQ water (without cells), continuous carbon grids (100 hexagonal mesh, EM Resolutions) were incubated in Poly-L-Lysine (PLL, Sigma, SLBZ7200) for 1 minute and 30 seconds, then the excess PLL was blotted away, followed by a 1-minute incubation with 5 µl Iron Sucrose or Iron citrate at a dilution of 1:2000. In contrast, 5 µl FCM was incubated for 1 minute at a dilution of 1:12,000 without prior PLL coating. All grids were then blotted with filter paper and air-dried.

For BF-TEM imaging of the iron formulations in cell culture medium, 1:2000 dilutions of IS, IC and FCM were incubated in RPMI medium with 10% FCS and 1% Pen-Strep for 24 h at 37 °C. After incubating continuous carbon grids (100 hexagonal mesh, EM Resolutions) in Poly-L-Lysine (PLL, Sigma, SLBZ7200) for 2 minutes and blotting the PLL excess away, 5 µl of the iron formulations were pipetted onto the grids and incubated for 1 minute at room temperature before blotting with filter paper. The grids were then washed quickly with MilliQ water to remove excess medium components, blotted with filter paper and air-dried.

For the comparison of the iron formulations in MilliQ water at pH∼7 *versus* enzyme-free artificial lysosomal fluid (ALF), the ALF mimicking endolysosomal conditions with a pH of ∼4.5 and a high salt concentration was prepared as described previously.^21,28^ Then, 1:10 dilutions of Iron Sucrose and Ferric Carboxymaltose were prepared in MilliQ or ALF and incubated at 37 °C for 24 h while rotating. Right after dilution (0 h), and then after 6 h and 24 h incubation, 5 µl of the dispersion was pipetted onto a continuous carbon grid (100 hexagonal mesh, EM Resolutions), incubated for 1 min at room temperature and washed twice with MilliQ water to remove the high salt concentration of the ALF that would lead to precipitation during drying. Then the grids were blotted with filter paper and air-dried.

BF-TEM imaging was performed for all grids with a Zeiss EM 900 microscope (Carl Zeiss Microscopy GmbH, Germany) at 80 kV and the Olympus iTEM 5.1 imaging software. Mean particle lengths were measured along their longest axis in TEM images using ImageJ 1.54b and statistics were calculated using Microsoft Excel 365.

### 11. High-Angle Annular Dark-Field (HAADF) Scanning Transmission Electron Microscopy (STEM) with Energy-Dispersive X-ray Spectroscopy (EDX)

For HAADF-STEM-EDX, the cell samples with particles were processed as for TEM with or without Turnbull’s blue staining, except that the ultrathin sections of 60-80 nm were placed onto Formvar-carbon coated copper grids (100 mesh, EM resolutions) after ultramicrotomy. The samples were then imaged with a Talos F200X TEM-STEM instrument (FEI, Super-X EDS) at an acceleration voltage of 200 kV and equal microscope settings for HAADF-STEM-EDX for all samples. The elemental maps were further analyzed using the Velox software (Version 3.0.0.815).

### 12. Computational Analysis of Endosome Size Differences

Instance segmentation of endosomes was performed following a semi-automated routine based on Segment Anything^29^ (vit_h model checkpoints) in a Napari^30^ interface (napari-segment-anything plugin) starting from expert prompts. Segmentation masks were then analyzed with a custom Python script (scikit-image^31^ v0.25), obtaining the area of each segmented organelle.

### 13. Measurement of Ferritin and Soluble Transferrin Receptor

To analyze ferritin and soluble transferrin receptor, 0.55 Mio cells per well were seeded in a 12-well plate (TPP), differentiated, polarized and treated with 1800 µM iron formulations for 0.75, 6, 14, 24 and 48 hours at 37°C. After treatment, the supernatant was collected, cells were washed twice with PBS, lysed in 500 µl lysis buffer (150 mM NaCl, 1% Triton-X, 50 mM Tris-HCl, pH 7.4), and supernatant as well as cell lysate were stored at -20 °C. Before analysis, the samples were thawed, and then the supernatant was centrifuged at 200 g for 5 min, while cell lysate was centrifuged at 14,000 g for 10 min. Only the liquid phase from each centrifugation was submitted to the Center for Laboratory Medicine for analysis by sequential chemiluminescent two-site immunoenzymatic sandwich assay. For this, samples were incubated with paramagnetic particles coated with monoclonal anti-ferritin (mouse) and goat anti-mouse IgG antibodies, along with a conjugate of goat anti-ferritin antibodies linked to alkaline phosphatase. The reagents were prepared in TRIS-buffered saline containing surfactant, bovine serum albumin, <0.1% sodium azide, and 0.1% ProClin™ 300.

### 14. Small-Angle X-ray Scattering (SAXS)

SAXS studies were carried out using a laboratory X-ray setup (Bruker Nanostar, Bruker AXS GmbH, Karlsruhe, Germany). Here, the K_α_-line of a micro-focused X-ray Cu source with an X-ray energy of 1.5406 Å was used. The beam was further passed through a 2D beam-shaping MONTEL mirror and then collimated using two 300 µm diameter pinholes, resulting in a beam diameter of approximately 400 µm at the sample position. An evacuated flight tube was placed between the sample and the detector to reduce absorption and air scattering. The scattered intensity of the materials placed in sealed quartz capillaries (1.5 mm, Hilgenberg GmbH) was recorded with a gaseous avalanche-based detector (VÅNTEC-2000, Bruker AXS). The sample-detector distance was set to 107 cm (as determined from measuring silver behenate powder as a standard) achieving a resolvable *q*-range of 0.07 ≤ *q* ≤ 2.3 nm^-1^. The scattering vector was given by

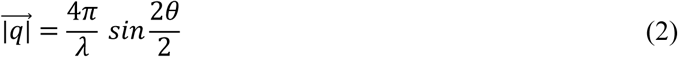

where 2θ is the scattering angle and λ is the wavelength of the X-ray source. The transmitted fraction of the beam was obtained from a homemade semi-transparent beam stop. The scattered intensity was extracted and then isotropic and azimuthally integrated over each *q*-value using the Bruker software DIFFRAC.EVA (Bruker AXS, version 4.1). The 1D scattering data was transmission corrected and then background subtracted from the scattering of the solvent and the quartz capillary using a home-built Python code.

Two different iron-carbohydrate complexes, IS and FCM, were diluted 1:50 and 1:100 in water and RPMI cell culture media (10% FCS, 1%PS), respectively. Then, SAXS measurements were conducted 1, 6 and 24 h after sample preparation with exposure times of either 1 h or 2 h.

### 15. Ultraviolet-Visible Spectroscopy (UV-VIS)

Absorbance measurements were performed using the UV-Vis Spectronometer (Cary 4000, Agilent Technologies). Iron formulations were prepared in Milli-Q water (IS/FCM), in RPMI cell culture media (IS/FCM RPMI) and RPMI cell culture medium supplemented with 10% FCS (IS/FCM RPMI (10% FCS)). After incubation of this solutions for 1 min or 1 h at room temperature, samples were analyzed in cuvettes (Plastibrand, PMMA, 759105) across a wavelength range of 200-800 nm. The baseline was corrected using Milli-Q water.

### 16. Immunofluorescence Staining of Rab 7, EEA1, LAMP1

For immunofluorescence staining PBS containing Ca²⁺ and Mg²⁺ was used for all steps. Polarized cells on 12 mm Ø coverslips were treated with 1800 µM iron formulations for 6 and 48 hours, washed once with PBS and fixed with 4% PFA for 20 minutes at room temperature. Cells were permeabilized with 0.1% Triton X-100 in PBS for 5 minutes at room temperature and subsequently washed three times with PBS for 5 minutes each. Coverslips were then removed from wells and placed on parafilm, where blocking was carried out by using 5% bovine serum albumin (BSA) in PBS for 30-60 minutes at room temperature. Mouse anti-Rab7 primary antibody (Cell Signaling, #95746) was diluted 1:200 in 1% BSA in PBS and applied overnight at 4 °C in a humidified chamber, followed by incubation with rabbit anti-EEA1 antibody (Abcam, #2900) diluted 1:900 in 1% BSA in PBS for 2 hours at room temperature. Mouse anti-Lamp1 antibody (Abcam, #25630) was diluted 1:100 in 1% BSA in PBS and incubated for 2 hours at room temperature. Secondary antibody goat anti-rabbit Alexa Fluor 488 (Invitrogen, #A11008) and goat anti-mouse Alexa Fluor 546 (Invitrogen, #A11030) were diluted 1:400 in 1% BSA in PBS and incubated for 1 hour at room temperature. Phalloidin 633 (Invitrogen, #A22284) was diluted 1:50 in 1% BSA in PBS and incubated for 30 minutes at room temperature. After and in between incubations, coverslips were always washed three times with PBS. Nuclear staining was performed using 1 µg/ml DAPI (Sigma, #D9542) for 10 minutes at room temperature, followed by two PBS washes and a final wash with Milli-Q water for a few seconds. Coverslips were then mounted using Mowiol and left to dry overnight at 37 °C before imaging. Antibodies were applied in a serial incubation manner, and all staining procedures were performed directly on coverslips placed on parafilm.

### 17. Western Blot Analysis of Rab7 and EEA1 Proteins

Human primary monocytes were seeded in 6-well plates (TPP) at a density of 1 Mio cells/well, differentiated, polarized and treated with iron formulations for 6 or 48 h as described above. Protein extraction was performed using RIPA buffer (Thermo/Pierce 89900) supplemented with HALT Protease Inhibitor Cocktail (Thermo 78429, 10 µl/ml and 0.5 mM PMSF). For EEA1 proteins analysis, the NuPAGE system was used, while for Rab7 protein analysis a standard SDS-PAGE protocol was followed. EEA1 proteins were separated using 4-12% Bis-Tris gels (1.5 mm thick, NP0335Box, Invitrogen). Samples were prepared by boiling in 4x LDS sample buffer containing 10 mM dithiothreitol (DTT) at 70 °C for 10 minutes, followed by centrifugation. For Rab7 analysis, proteins were separated using a 12% acrylamide gel prepared according to standard protocols. Samples were mixed with 6× SDS loading buffer containing 10 mM DTT, boiled at 95 °C for 5 minutes, and briefly centrifuged. Electrophoresis for EEA1 analysis was performed using MOPS running buffer, supplemented with the recommended antioxidant for the NuPAGE Bis-Tris Gel System (Invitrogen). MOPS running buffer (20×) was prepared by dissolving 104.6 g MOPS (Sigma M1254), 60.6 g Tris base (Sigma 14880), 10 g SDS, and 3 g EDTA (Sigma 3620) in water to a final volume of 500 ml, adjusting the pH to 7.7. For Rab7 proteins, Tris-Glycine-SDS running buffer was utilized. A total of 15 µg of protein per condition was loaded per well, alongside 10 µl of a PageRuler Protein Ladder (Thermo). The gel was run at 80 V for 20 minutes, followed by 120 V until the dye front exited the gel or optimal separation of high-molecular-weight proteins was achieved. For Western blotting, polyvinylidene difluoride (PVDF) membranes (Invirolon, Invitrogen LC2005) were pre-soaked in 100% methanol prior to transfer. EEA1 samples were transferred onto these membranes overnight at 4 °C using a Bio-Rad wet blot system, with additional cooling provided by an ice block. Rab7 proteins were also transferred onto a PVDF membrane using the iBlot system (Thermo Fisher Scientific). Transfer buffer (20×) was prepared by dissolving 10.2 g Bicine, 13.1 g Bis-Tris (free base), and 0.75 g EDTA in water to a final volume of 125 ml, with a final pH of 7.2. Working transfer buffer (1 l) was prepared by diluting 50 ml of 20× transfer buffer with 200 ml methanol (100 ml per gel) and 749 or 849 ml Milli-Q water, including 1 ml of NuPage antioxidant. Following transfer, membranes were blocked for 1 hour at room temperature in either 5% Quick Blocker (Chemicon, 2080) for EEA1 or 5% BSA in PBST (PBS containing 0.05% Tween-20) for Rab7. Membranes were then incubated overnight at 4 °C with primary antibodies against EEA1 (1:1000, rabbit, ab2900, Abcam) or Rab7 (1:1000, mouse, 95746, Cell Signaling) in 2.5% BSA/PBST. GAPDH-HRP (Cell Signaling 3683S, 1:1000) was used as a loading control. After three 5-minute washes in PBS at room temperature, membranes were incubated with horseradish peroxidase (HRP)-conjugated anti-rabbit or anti-mouse secondary antibody (1:10,000, RPN4301 GE Healthcare, AP124P Chemicon) in 2.5% BSA/PBST for 1 hour at room temperature. After three additional washes with PBS, chemiluminescent detection was performed using the Pierce/Thermo SuperSignal West Pico substrate (Thermo #34080). Membranes were imaged for chemiluminescence, and a separate image was taken to document the molecular weight marker (BioRad ChemiDoc MP).

### 18. Proteomics Data Analysis

We analyzed the mass-spectrometry (MS) proteomics data from Bossart *et al.* quantifying protein levels in macrophages treated with IS or FCM for 6 h as well as non-treated macrophages.^15^ We used the DEP Bioconductor package (version 1.26.0)^32^ to run the differential expression analysis.

We first filtered out contaminants and entries matching the reversed part of the decoy database. Next, we filtered out proteins that were identified in all replicates of at least one condition. We then background corrected and normalized the expression data using the variance stabilizing transformation, followed by missing value imputation using a normal distribution with a negative shift of 1.4 standard deviations from the mean and a width of 0.4 standard deviations of the measured values. Principal component analysis of the normalized data showed a donor effect in the proteomic profiles (Suppl. Fig.6f). Therefore, we included the donor information in the design to test for the effect of the treatment while controlling for donor.

The code for the differential analysis of the proteomics data can be found at: https://github.com/rriupu/iron_carbohydrate_ms_data_analysis.git.

### 19. Statistical Analysis

Graph generation and statistical analysis were performed using GraphPad Prism unless stated otherwise in the respective methods sections. Unless otherwise noted, results are represented as the mean ± standard error of the mean (S.E.M.). Comparisons between two independent groups were conducted using an unpaired *t* test. We used the following convention to indicate significance with asterisks: not significant (ns), (P > 0.05), *(0.05 > P > 0.01), **(0.01 > P > 0.001).

