## Supplementary Data for "Understanding the kinetics of macrophage uptake and the metabolic fate of iron-carbohydrate complexes used for iron deficiency anemia treatment"

### Material and Methods

#### 1. Dosimetry assessment

To investigate the effect of media volume on cellular iron formulation uptake, a dosimetry experiment was conducted using THP-1 cells. $2.3 \times 10^5$ cells were seeded with 200 nM PMA in 24-well plates (TPP) and incubated for 3 days in RPMI-1640 (Sigma-Aldrich) culture medium (supplemented with 10% fetal calf serum (FCS), 1% penicillin-streptomycin (PS) and 2 mM L-Glut) at 37 °C with 5% $CO_2$. Cells were treated with either 200 µl or 580 µl per well of 1800 µM iron formulation and incubated for 0.75/6/24 hours at 37 °C with 5% $CO_2$. After treatment, the medium was removed, and cells were washed twice with 1 ml PBS before fixation with 500 µl of 4% PFA in PBS for 20 minutes at room temperature in a fume hood. Perls Prussian blue and DAB staining were then performed as previously described. All experiments were performed in triplicates to ensure reproducibility.

#### 2. Fluorescent bead uptake

##### 1. Impact of ICC Sugar Shell on Fluorescent Bead Uptake

Human primary monocytes cells were seeded at a density of $2 \times 10^5$ cells per well in 8-well plates (Ibidi, Cat. No. 80826) in a 30 µl of RPMI media drop. After 6 h, the cells have settled down and the well was filled with fresh differentiation media. Following differentiation and polarization, cells were pretreated using a concentration of 0.5, 5, and 10 g/ml of Sucrose (CSL Vifor, Switzerland) or Carboxymaltose (CSL Vifor, Switzerland) dissolved in RPMI medium at 37 °C for 30 minutes. After pre-treatment, 300 µl of Yellow Sphero-beads (Spherotech, FP-00552-2) or $SiO_2$ FITC beads (KRISS) were added to the wells. The plates were incubated for 1.5 hours at 37 °C in the dark. Cells were washed three times with PBS and fixed by 4% PFA for 20 minutes at room temperature. Following fixation, the cells were incubated in a 1:1000 dilution of DAPI (4',6-diamidino-2-phenylindole) in PBS for 15 minutes in the dark at room temperature, before being washed three times with PBS to remove excess DAPI. Confocal laser scanning microscopy (CLSM) was employed to visualize the cells and bead interactions with a 40x objective, capturing at least 10 Z-stacks with steps of approximately 1.2 µm. The following settings were applied for imaging:

| | **Dapi** | **Yellow Sphero-beads** | **$SiO_2$ FITC beads** |
|---|---|---|---|
| Gain | 485 | 670 | 739 |
| Dig. Offset | 0 | 0 | 0 |
| Dig. Gain | 1.0 | 1.0 | 1.0 |
| Pinhole | 89.7 | 89.7 | 89.7 |

2. Effects of Amantadine and Cytochalasin D on Fluorescent Bead Uptake

After differentiation and polarization, cells were pretreated with 0.5 mg/ml Amatadine and 0.5 µM Cytochalasin D for 1h, followed by incubation with 100 µl of Yellow Sphero-beads (Spherotech, FP-00552-2) or $SiO_2$ FITC beads (KRISS) containing the appropriate dilution of inhibitor and nanoparticle. The plates were incubated for 1.5 hours at 37 °C in the dark. Cells were washed three times with PBS and fixed by 4% PFA for 20 minutes at room temperature. Following fixation, the cells were incubated in a 1 µg/ml dilution of DAPI (4',6-diamidino-2-phenylindole) in PBS for 10 minutes in the dark at room temperature, before being washed three times with PBS to remove excess DAPI. Confocal laser scanning microscopy (CLSM) was employed to visualize the cells and bead interactions with a 40x objective, capturing at least 10 Z-stacks with steps of approximately 1.2 µm. The following settings were applied for imaging: Laser: Dapi: 0.5%; FITC: 1.5; Yellow: 1.2%

| | **Dapi** | **Yellow Sphero-beads** | **$SiO_2$ FITC beads** |
|---|---|---|---|
| Gain | 750 | 650 | 700 |
| Dig. Offset | 20 | 20 | 20 |
| Dig. Gain | 1.0 | 1.0 | 1.0 |
| Pinhole | 0.81AU | 0.66AU | 0.66AU |

3. Image analysis

Microscopy images were analyzed using Fiji (ImageJ, NIH). For each image stack, a maximum intensity projection was generated. The resulting images were then split into individual color channels to isolate the signal of interest. Quantification of the total area in the relevant fluorescence channel was performed using the “Analyze Particles” plugin, following appropriate thresholding. To account for variations in cell density, the total area measured per field of view was normalized to the number of nuclei present in the same field.

#### 4. MTS Assay of endocytic inhibitors

Human monocyte cells were seeded in 96-well plates (TPP) at a density of $4 \times 10^4$ cells per well in 100 µl of medium, differentiated, polarized, and pretreated for 1 hour with 0.1, 0.3, or 0.5 mg/ml Amantadine (Sigma, A1260), 0.1, 0.3, or 0.5 µM Cytochalasin D (Sigma, C8273), and 0.1% DMSO and Milli-Q water as controls. Following preincubation, the inhibitors were co-incubated with 1800 µM iron for 6 hours. After medium was completely removed, cell viability was assessed using an MTS assay kit (CellTiter 96 AQ; Promega, G3581). Therefore, RPMI medium without phenol red (Sigma, R7509) was mixed with MTS reagent and a total of 120 µl of this mixture was added to each well. Plates were incubated for 60 minutes at 37°C with 5% $CO_2$, and optical density was measured at 490 nm using a microplate reader (BioTek, Synergy H1). All experiments were performed in triplicates to ensure reproducibility.

### Supplementary Figures

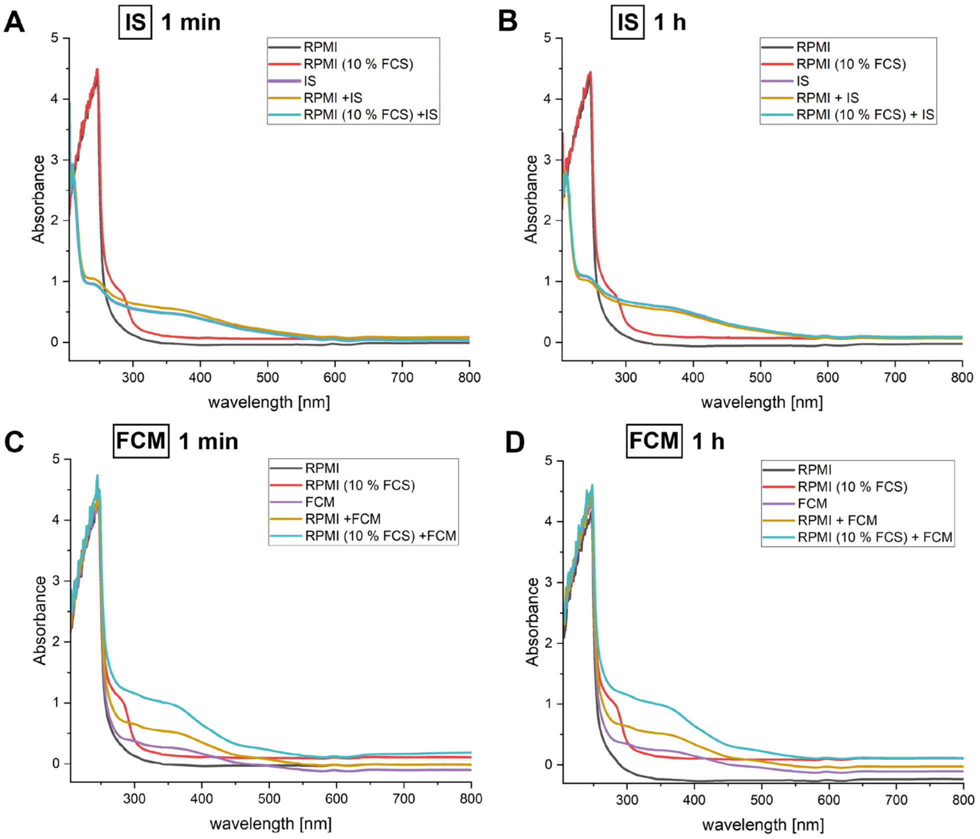


**Supplementary Figure 1 to Main Figure 1: Characterization of iron carbohydrate complexes (ICCs) in cell culture medium. A-D)** UV-VIS measurements of IS and FCM incubated for 1 min or 1 h with either RPMI (without FCS) or complete cell culture medium (RPMI + 10% FCS + 1% Pen-Strep). The curves of IS (purple) and IS in RPMI (10% FCS, 1% Pen-Strep, turquoise) are overlapping in **A-B)**. IS: iron sucrose, FCM: ferric carboxymaltose. Absorbance: arbitrary units.

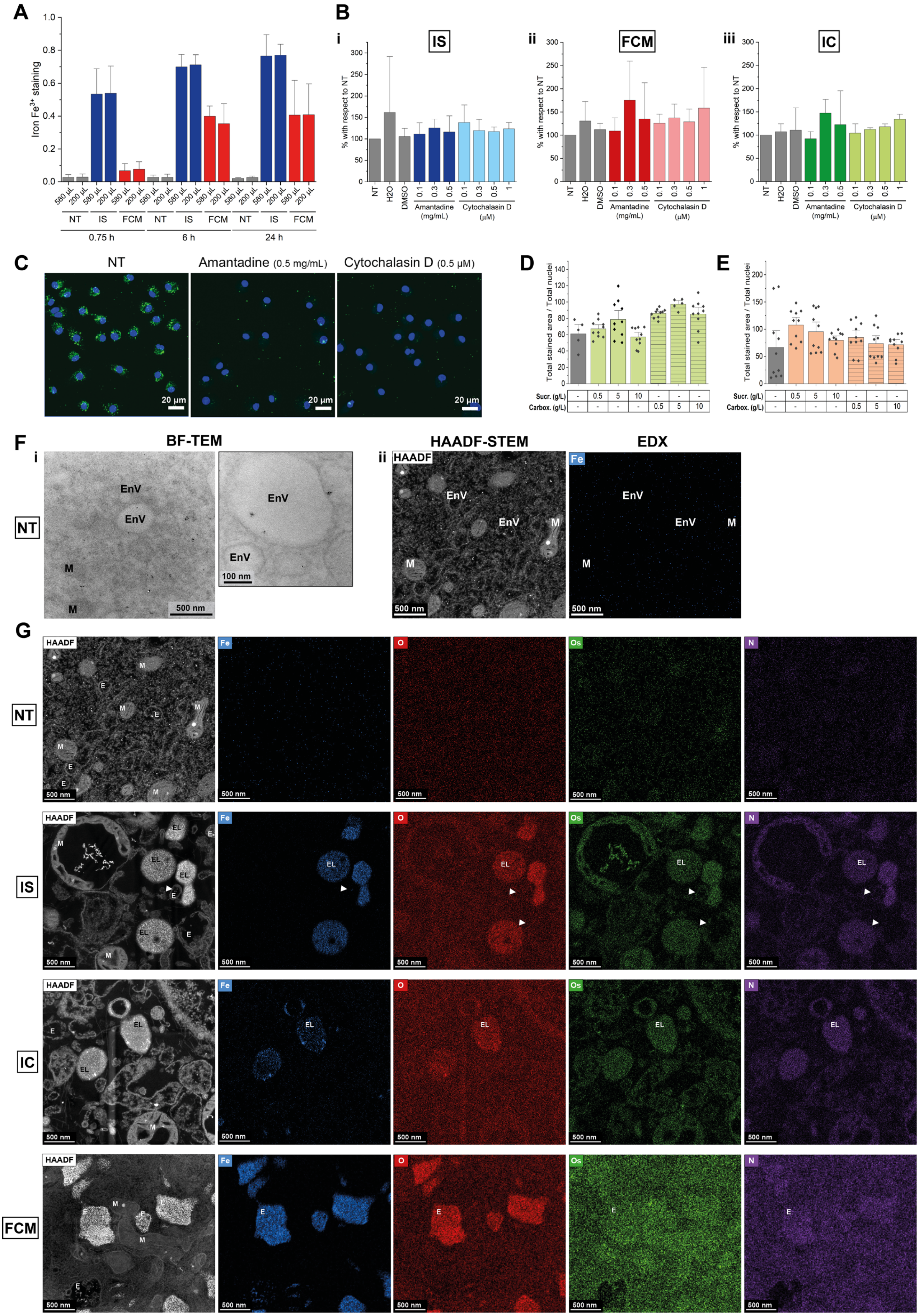

A
Iron $Fe^{3+}$ staining
580 µL
200 µL
NT
IS
FCM
0.75 h
6 h
24 h
B
i
IS
ii
FCM
iii
IC
% with respect to NT
NT
H2O
DMSO
Amantadine (mg/mL)
Cytochalasin D (µM)
C
NT
Amantadine (0.5 mg/mL)
Cytochalasin D (0.5 µM)
20 µm
D
E
Total stained area / Total nuclei
Sucr. (g/L)
Carbox. (g/L)
F
i
ii
BF-TEM
HAADF-STEM
EDX
HAADF
Fe
EnV
M
500 nm
100 nm
G
HAADF
Fe
O
Os
N
NT
IS
IC
FCM
EL
E
M
500 nm

**Supplementary Figure 2 to Main Figure 2: Dynamics of cell uptake, endocytosis and role of carbohydrate ligand. A)** Dosimetry assessment of IS and FCM in THP-1 derived macrophages. The cells were treated over time (45 min, 6 h and 24 h) with either 200 µL or 580 µL of the ICCs solution. They were then fixed, stained with Prussian blue and DAB, and the $Fe^{3+}$ staining was quantified with ImageJ. The bar graphs represent the mean + S.E.M. of triplicates. NT: non-treated. **B)** The cytotoxicity of the indicated concentrations of Amantadine and Cytochalasin D was assessed with an MTS assay in primary human macrophages. The bar graphs represent the mean + S.E.M. of triplicates with three different donors. IC: labile $Fe^{3+}$ iron control iron citrate; $H_2O$: water; DMSO: dimethyl sulfoxide as carrier control of the inhibitors. **C)** The inhibitory activity of Amantadine and Cytochalasin D on endocytosis was confirmed with polystyrene fluorescent beads. As with the ICCs in Fig.2, primary human macrophages were pre-treated with the inhibitors for 30 min, then the beads were added. The cells were processed for microscopy and imaged with a confocal microscope. Shown are representative images of the highest concentrations of Amantadine (0.5 mg/mL) and Cytochalasin D (0.5 µM). Color code: DAPI nuclear staining (blue), fluorescent beads (green). Scale bar: 20 µm. **D-E)** To assess the effect of the carbohydrate ligands on the uptake of non-ICCs nanoparticles, **D)** silica nanoparticles (20 nm) and **E)** Sphero-beads (70 nm) were used. The macrophages were pretreated for 30 min with the indicated concentrations of sucrose or carboxymaltose. Afterwards, silica nanoparticles (**D**) or Sphero-beads (**E**) were added at a concentration of 100 µg/ml, respectively. After 6 h of incubation in the presence of the ligands, the cells were washed, fixed and processed for fluorescence microscopy. Uptake was calculated with ImageJ, by quantifying the total area of the fluorescence channel corresponding to the nanoparticles (Analyze Particles plugin). This value was then normalized to the number of nuclei per field of view. Shown are the bar graphs of a duplicate + S.E.M. **F)** Representative images of bright field (BF)-TEM (**i**, with zoom inset) and scanning TEM (STEM) coupled with high-angle annular dark-field imaging (**ii**, HAADF) of non-treated (NT) primary macrophages after 24 h as control, showing no iron NPs inside endocytic vesicles (EnV) as expected and a very low, disperse iron (Fe) background signal in the cell in in elemental analysis *via* energy-dispersive X-ray spectroscopy (**ii**, EDX). M: mitochondria. Scale bars: BF-TEM: 500 nm, 100 nm; HAADF-STEM-EDX: 500 nm. **G)** Representative images of HAADF-STEM imaging (left) and elemental analysis after 24 h *via* EDX showing that based on the analysis with Turnbull's staining in Fig.5 and Suppl. Fig.5, also in the unstained EM and EDX images, the EnV with low and diffusely distributed osmium (Os) and nitrogen (N) signals close to the cell background can be attributed as endosomes (E) with no digestive enzymes in contrast to the endolysosomes (EL) with a high concentration of digestive enzymes and thus strong Os and N signals. IS NPs and IC precipitates with localized iron (Fe) and oxygen (O) signals are already in EL at 24 h, while FCM NPs are still in E. Scale bars: 500 nm.

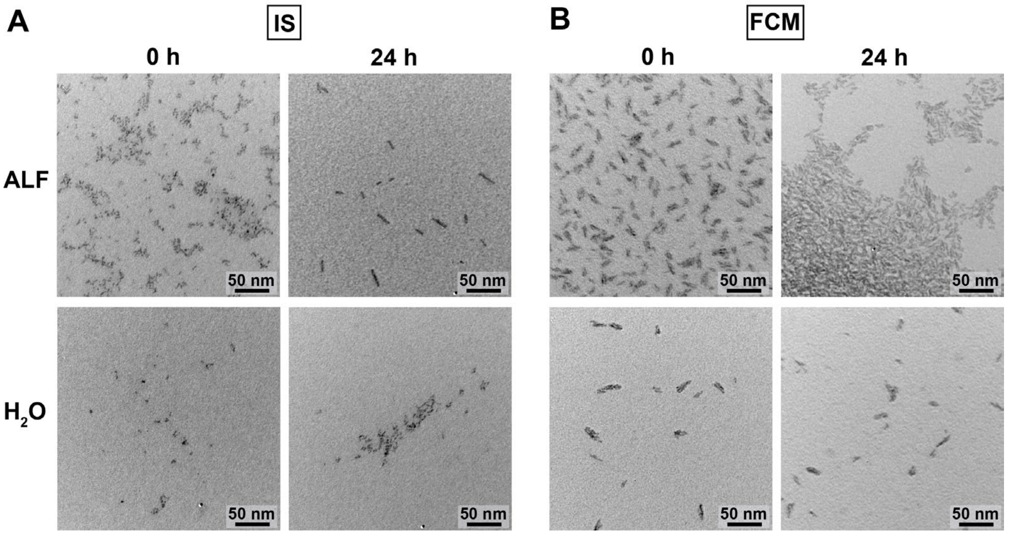


**Supplementary Figure 3 to Main Figure 3: *In vitro* pharmacokinetic profiling of ICCs. A-B)** Representative BF-TEM images of IS and FCM NPs incubated for 24 h rotating at 37 °C in artificial lysosomal fluid (ALF, pH 4.5) or water. **A)** Only elongated clusters of IS NPs are left after 24 h in ALF, while the individual single spherical NPs are disappearing *via* dissolution to iron ions invisible with BF-TEM. In water ($H_2O$), the single spherical NPs remain visible and in similar abundance over time. **B)** Ellipsoidal FCM clusters retain their shape, size and abundance in endolysosome-mimicking ALF over time, while showing smaller inter-cluster distances, likely due to charge screening of the ligand charges by the salt ions of the ALF. In contrast, the FCM clusters retain a similar size and inter-cluster distance in water over time. Scale bars: 50 nm.

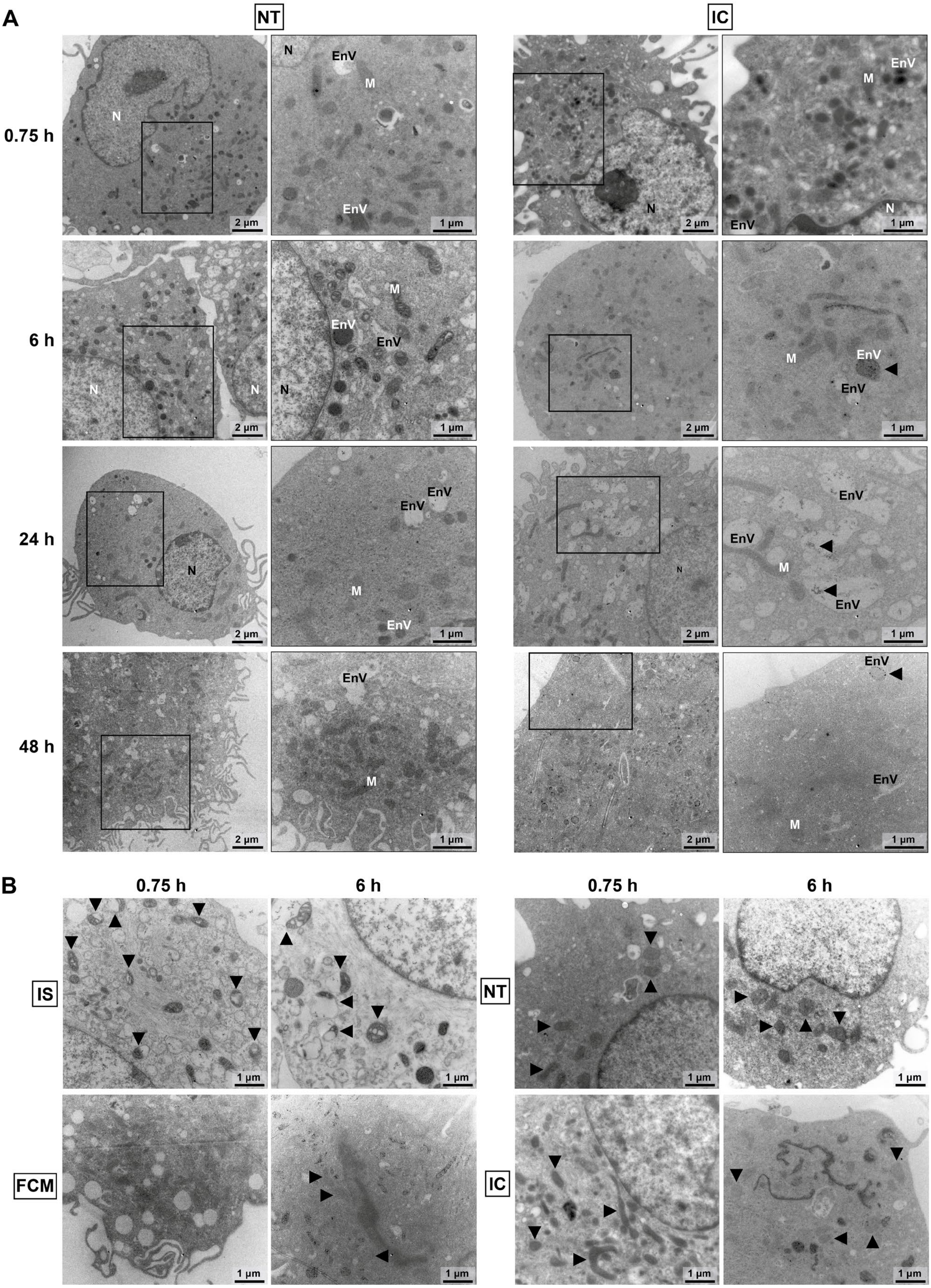
A
NT
IC
0.75 h
6 h
24 h
48 h
N
EnV
M
2 µm
1 µm
B
0.75 h
6 h
IS
FCM
NT
IC

**Supplementary Figure 4 to Main Figure 4: Intracellular trafficking. A)** Representative BF-TEM images showing the controls of non-treated (NT) primary macrophages and iron citrate (IC) as $Fe^{3+}$ control over time at different magnifications (black squares). In endocytic vesicles (EnV) of IC-treated cells, precipitates are visible (black arrows) as seen in main Fig.1 in water and medium. N: nucleus, M: mitochondria. Scale bars: 2 µm, 1 µm. **B)** At early time points, IS NPs led to swollen mitochondria (black arrows) with obvious cavities visible in BF-TEM, indicative of oxidative stress, which was reversible over time and not observed in non-treated cells (NT) or FCM/IC-treated primary macrophages (black arrows). Scale bars: 1 µm.

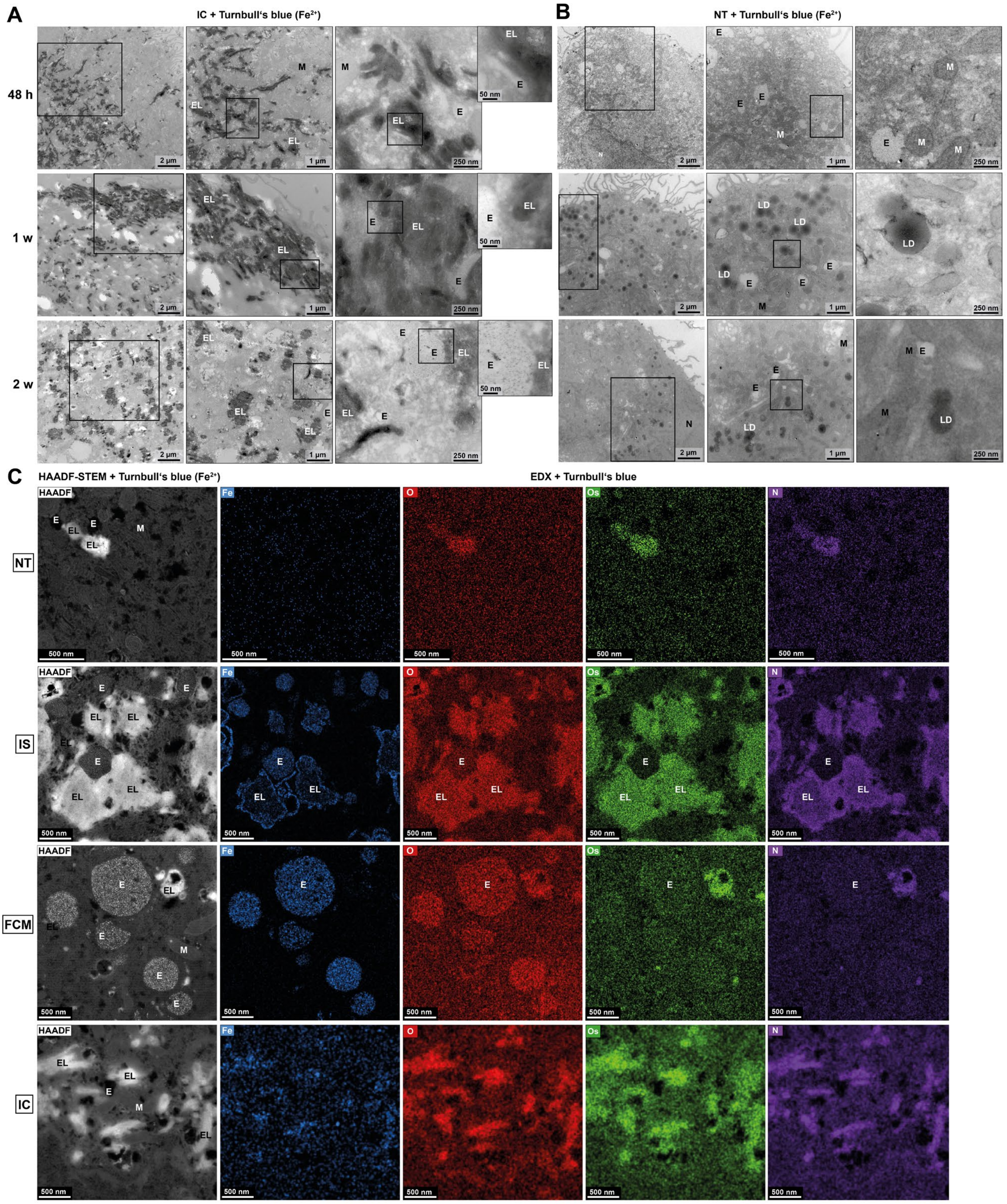


**Supplementary Figure 5 to Main Figure 5: *The Hamster Effect*. A-B)** Representative images of BF-TEM combined with Turnbull's blue staining enhanced with DAB to visualize the digestion of $Fe^{3+}$ in **A)** the $Fe^{3+}$ control iron citrate (IC) and **B)** non-treated (NT) primary macrophages to $Fe^{2+}$ in endolysosomes (EL) at 48 h, 1 week (1 w) and 2 weeks (2 w) after uptake at different magnifications (black squares). **A)** For IC, many strongly Turnbull's-stained EL (dark structures) were visible at 48 h, which decreased in abundance in 2 weeks, as well as much smaller endosomes (E) without iron digestion and thus Turnbull's staining. **B)** Non-treated primary macrophages survived for 2 weeks on the iron in the medium, but showed signs of stress in the form of autophagy-related lipid droplets (LD). N: Nucleus, M: Mitochondria. Scale bars: 2 µm, 1 µm, 250 nm, 50 nm. **C)** Combination of representative HAADF-STEM (left) images with Turnbull's staining (enhanced with DAB) and EDX at 48 h revealing that the Turnbull's-stained and thus $Fe^{2+}$-positive EnV are EL with characteristic osmium (Os) and nitrogen (N) signals from the endosomal digestive enzymes, and with iron (Fe) and oxygen (O) signals from still undigested IS/FCM NPs inside the EL. FCM NP-filled EnV are devoid of OS and N signals, indicating that they are E without digestive enzymes. Upon IC-treatment, localized Fe and O signals of IC precipitates in EL with Os and N signal are visible, while in non-treated cells the Fe background signal is diffusely distributed over the entire cell, with O, Os and signals localizing to EL as expected from their high digestive enzyme and salt concentrations. Scale bars: 500 nm.

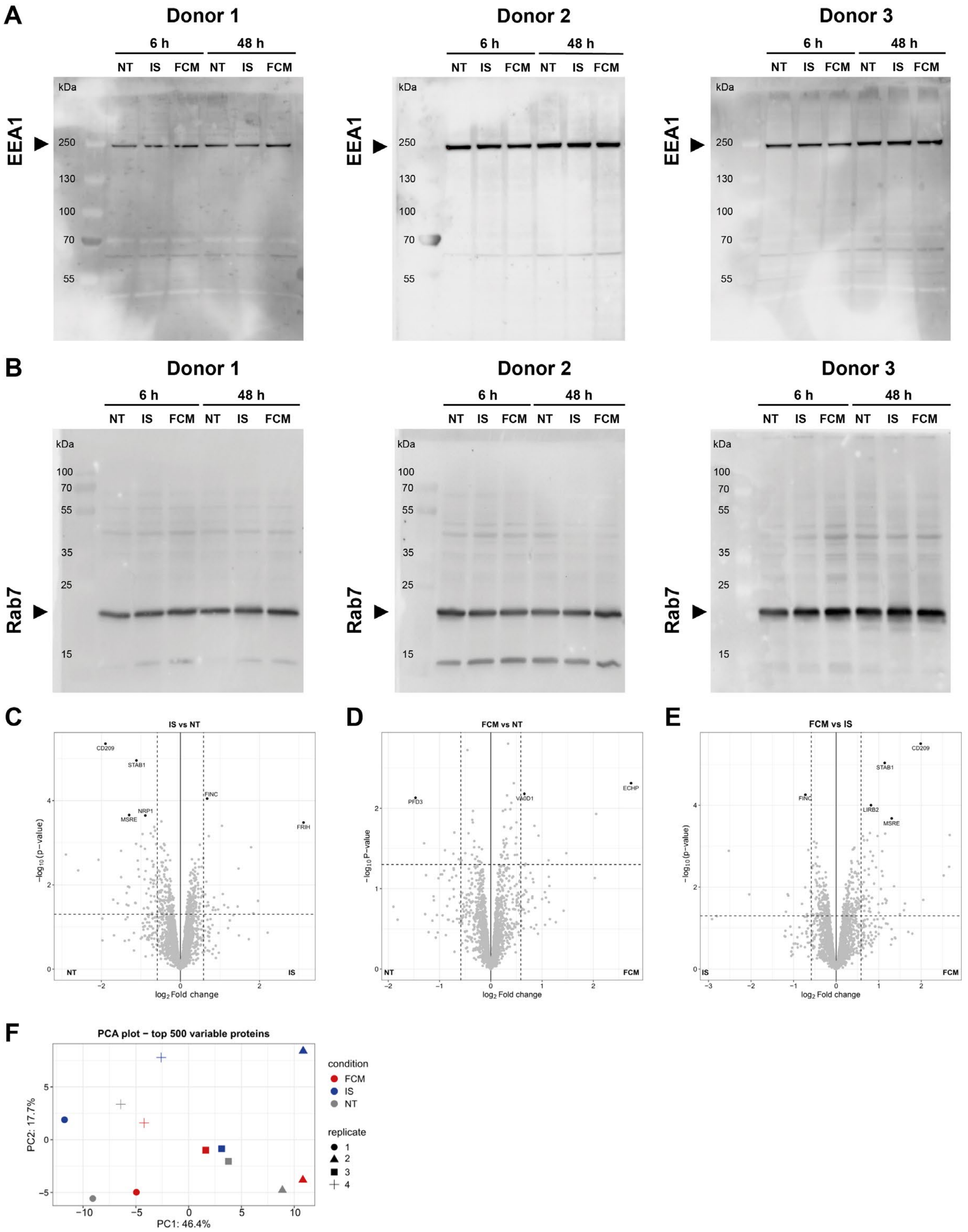


**Supplementary Figure 6 to Main Figure 6: Mechanism of action of ICCs in macrophages. A)** Western blot analysis of EEA1 protein levels (black arrow) in samples from three different donors treated with either IS or FCM for 6 h and 48 h. Each blot represents an individual donor. **B)** Western blot analysis of Rab7 protein levels (black arrow) in the same three donors. **C-E)** Volcano plots of proteomics showing the differentially expressed proteins in <condition1> *versus* <condition2> (link to database in Main Materials and Methods). The black dots show the proteins passing the adjusted p-value threshold after multiple testing correction. **F)**

PCA: Principal Components 1 (x-axis) and 2 (y-axis) obtained when using the top 500 most variable proteins across all samples.
